# Environmental palaeogenomic reconstruction of an Ice Age algal population

**DOI:** 10.1101/2020.04.10.035535

**Authors:** Youri Lammers, Peter D. Heintzman, Inger Greve Alsos

## Abstract

Palaeogenomics has greatly increased our knowledge of past evolutionary and ecological change, but has been restricted to the study of species that preserve as fossils. Here we show the potential of shotgun metagenomics to reveal population genomic information for a taxon that does not preserve in the body fossil record, the algae *Nannochloropsis*. We shotgun sequenced two lake sediment samples dated to the Last Glacial Maximum and identified *N. limnetica* as the dominant taxon. We then reconstructed full chloroplast and mitochondrial genomes to explore within-lake population genomic variation. This revealed at least two major haplogroups for each organellar genome, which could be assigned to known varieties of *N. limnetica*. The approach presented here demonstrates the utility of lake sedimentary ancient DNA (*sed*aDNA) for population genomic analysis, thereby opening the door to environmental palaeogenomics, which will unlock the full potential of *sed*aDNA.

## Introduction

Palaeogenomics, the genomic-scale application of ancient DNA, is revolutionizing our understanding of past evolutionary and ecological processes, including population dynamics, hybridization, extinction, and the effects of drivers of change ^1–4^. Despite extensive application to, and innovations using, body fossils ^5–7^, its use on another major source of ancient DNA - the environment - has been almost entirely limited to inferring the presence or absence of taxa through time ^8–14^. However, a nuanced understanding of ecological and evolutionary dynamics requires population genomic information. The direct recovery of this information from cave sediment has recently been shown ^12^, but - to our knowledge - has not yet been demonstrated for lake sediments.

Lake sediments provide an ideal source of sedimentary ancient DNA (*sed*aDNA) that originates from both the catchment and the lake itself, as well as providing a stable environment required for optimal aDNA preservation ^15,16^. As a result, lake *sed*aDNA has been used to infer the taxonomic composition of past communities ^16,17^, regardless of whether those taxa preserve in the body fossil record. The most commonly applied method is DNA metabarcoding, which allows for the targeting of particular groups of organisms ^18,19^. However, the ability to confidently identify barcodes is constrained by the completeness of appropriate reference databases, and the length and variability of the barcode targeted. Short barcodes are necessarily targeted for fragmented aDNA ^20^, which can therefore impede species-level identification. An alternative approach is shotgun metagenomics, which is non-targeting and preserves aDNA damage patterns that, in contrast to metabarcoding, allows for authentic aDNA to be distinguished from modern contamination ^16,21–23^. For palaeogenomic reconstruction however, either deep shotgun sequencing or target enrichment of *sed*aDNA is required, which allows for robust species-level identification ^12,14,24^, as well as the potential exploration of population genomic variation.

Andøya, an island located off the coast of northwest Norway, was partially unglaciated during the Last Glacial Maximum (LGM, Figure 1) and has therefore been a focus of palaeoecological studies ^25^, especially for its potential as a cryptic northern refugium ^26–28^. Studies focussing on sediment cores from three lakes (Endletvatn, Nedre Æråsvatnet, Øvre Æråsvatnet)^26,29–35^ have reported the presence of an Arctic community during the LGM, which includes taxa such as grasses (Poaceae), crucifers (Brassicaceae), and poppy (*Papaver*), along with bones of the Little Auk (*Alle alle*). Furthermore, recent geochemical and DNA metabarcoding analyses indicate the presence, and an inferred high abundance, of the algae *Nannochloropsis* in LGM sediments from Andøya ^35^.

**Figure 1:**
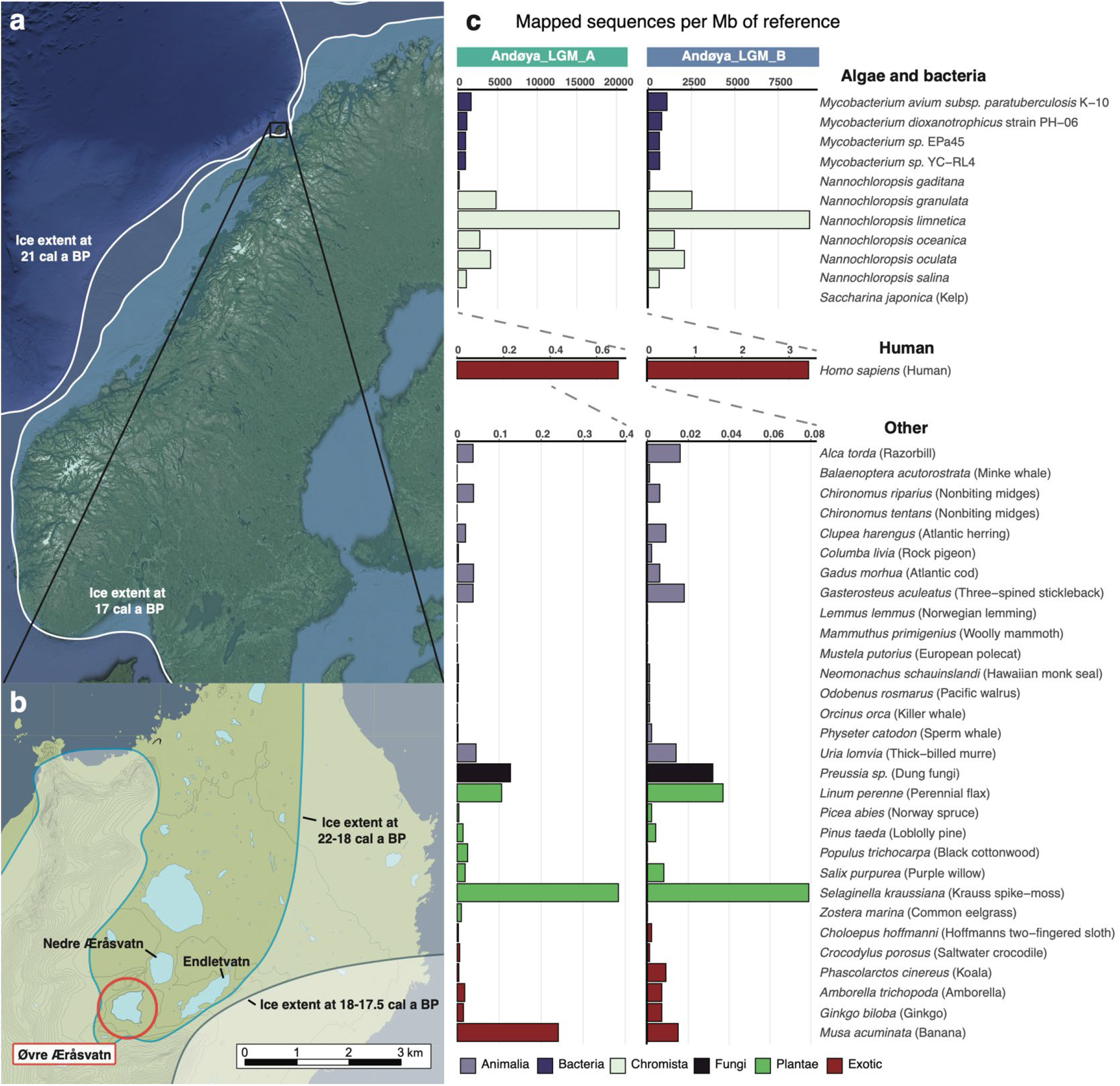
(a, b) Location of Lake Øvre Æråsvatnet (circled in red, panel b) in the ice-free refugium of Andøya in northwest Norway. The regional ice extent for Scandinavia in panel a has been plotted for 22 (outer) and 17 (inner) kcal yr BP and is based on Hughes et al.^78^. The local ice extent in panel B is plotted for 22-18 and 18-17.5 kcal yr BP and is based on Vorren et al. ^25^. (c) Taxonomic composition of the LGM Andøya sediment samples, based on alignment to a reference panel of 42 eukaryotic or bacterial nuclear genomes. For readability, the algal, bacterial, and human results are plotted separately with differing y-axis scales.

*Nannochloropsis* is a genus of single-celled microalgae of the Eustigmatophyceae. All species have high lipid contents and are therefore of interest as a potential source of biofuels ^36,37^. As a result of this economic interest, the organellar and nuclear genomes have been sequenced for six of the eight described species (Supplementary Table S1) ^37–40^. All species are known from marine environments, with the exception of *N. limnetica*, which is known from freshwater and brackish habitats, and comprises five varieties ^41–43^. The genus has a cosmopolitan distribution, with the marine species being reported from most oceans ^44–46^, whereas the freshwater/brackish *N. limnetica* is known from lakes in Europe ^41^, Asia ^42^, North America ^43,47^, and Antarctica ^48^. Species-level identification of *Nannochloropsis* from water is problematic, due to its small size (2 to 6 μm in diameter ^43,46^) and, in contrast to diatoms ^49^, lack of diagnostic morphological structures ^47,50^. Reliable species identification is however possible with short genetic markers ^43,47,51^. In sediments, *Nannochloropsis* has not been reported from macrofossil and pollen/spore profiles, and may therefore only be identifiable using *sed*aDNA ^10,35,52^.

In this study, we shotgun sequenced two broadly contemporaneous LGM lake sediment layers from Andøya that a previous metabarcoding study had shown to contain *Nannochloropsis* ^35^. The depth of our shotgun metagenomic data, together with the availability of a reference genome panel, allowed us to demonstrate that *N. limnetica* dominates the identifiable taxonomic profile. Through reconstruction of complete chloroplast and mitochondrial genomes, we show that at least two variants of *N. limnetica* are represented. We thus demonstrate, and to the best of our knowledge for the first time, that it is possible to estimate past population genomic diversity both from total *sed*aDNA and from a taxon not preserved in the body fossil record.

## Results

### Metagenomic analysis and species-level determination of *Nannochloropsis*

We shotgun sequenced two LGM samples, dated to 17,700 (range: 20,200-16,500) and 19,500 (20,040-19,000) calibrated years before present (cal yr BP), to generate 133-224 million paired-end reads, of which we retained 53-127 million sequences after filtering (Supplementary Table S2). We first sought to identify the broad metagenomic profiles of the samples and the species-level identification of *Nannochloropsis* from Lake Øvre Æråsvatnet.

First, for each sample, we compared two non-overlapping one-million sequence subsets of the filtered data to the NCBI nucleotide database. The taxonomic overlap between the two one million read subsets was 88-93% within each sample, demonstrating that our subsets are internally consistent. We then merged the two subsets from each sample, which resulted in the identification of 29,500-32,700 sequences (Table 1). The majority of the identified sequences were bacterial, with 21-26% identified as *Mycobacterium*, although the majority of these sequences could not be identified to a specific strain. Within the eukaryotes, *Nannochloropsis* constituted ~20% of the assigned sequences in both samples, with ~33% of these identified as *N. limnetica* (Table 1; Supplementary Figure S1; Supplementary Table S3).

**Table 1:** Summary of the best represented taxa (>200 identified sequences) detected in the metagenomic analysis. N=Number of identified sequences, I=Percentage of identified sequences, A=Percentage of all sequences included in the metagenomic analysis.

|  | Andøya_LGM_A |  |  | Andøya_LGM_B |  |  |
| --- | --- | --- | --- | --- | --- | --- |
|  | N | I | A | N | I | A |
| Bacteria | 18,852 | 63.93 | 0.94 | 21,873 | 66.85 | 1.09 |
| *Mycobacterium* | 6268 | 21.26 | 0.31 | 8535 | 26.08 | 0.43 |
| *M. avium* complex | 444 | 1.51 | 0.02 | 635 | 1.94 | 0.03 |
| *M. dioxanotrophicus* | 0 | 0 | 0 | 229 | 0.7 | 0.01 |
| *M. sp.* EPa45 | 0 | 0 | 0 | 306 | 0.94 | 0.02 |
| *M. sp.* YC-RL4 | 0 | 0 | 0 | 207 | 0.63 | 0.01 |
| *Pseudomonas* | 920 | 3.12 | 0.05 | 904 | 2.76 | 0.05 |
| Eukaryota | 9333 | 31.65 | 0.47 | 9563 | 29.23 | 0.48 |
| *Nannochloropsis* | 5913 | 20.05 | 0.3 | 6179 | 18.88 | 0.31 |
| *N. limnetica* | 2223 | 7.54 | 0.11 | 2272 | 6.94 | 0.11 |
| Other | 1303 | 4.42 | 0.07 | 1284 | 3.92 | 0.06 |
| Total | 29,488 | 100 | 1.47 | 32,720 | 100 | 1.64 |

To further investigate the metagenomic profile of the samples, we aligned all filtered sequences to each nuclear genome within a reference panel derived from four *Mycobacterium* strains and 38 eukaryotes, with the latter either exotic (implausible) or non-exotic (plausible) to LGM Andøya (Supplementary Table S4). We mapped 310,000-680,000 sequences to the *N. limnetica* genome, which translates to 9.3-20.3 thousand sequences per megabase (kseq/Mb), and a nuclear genomic coverage of 0.48-1.13x. We observed a far lower relative mapping frequency to all other *Nannochloropsis* nuclear genomes (up to 2.5-4.8 kseq/Mb). If we consider sequences that are only mappable to a single genome, then the relative mapping frequency falls to 7.4-17 kseq/Mb for *N. limnetica* and up to 0.6-1.3 kseq/Mb for all other *Nannochloropsis* genomes. The most abundant non-*Nannochloropsis* eukaryotic taxon in the sequence data was human, with 2-11 thousand sequences mapped (0.7-3.4 seq/Mb). As expected from the metagenomic analyses, the next most abundant group is *Mycobacterium*. As the relative mapping frequency was consistent across all four strains, based on both all sequences aligned (up to 1.1-1.7 kseq/Mb) and only retaining sequences unique to a strain (up to 0.6-0.9 kseq/Mb), we infer that the *Mycobacterium* strain or strains present in LGM Andøya are not closely related to any that have been sequenced to date (Figure 1; Supplementary Table S4). The relative frequencies of sequences mapping to plausible and implausible eukaryotic genomes are comparable for non-*Nannochloropsis* taxa. Based on both raw counts and those corrected for genome size, these analyses therefore indicated that *N. limnetica* is the best represented taxon in the panel (Figure 1; Supplementary Table S4).

We sought to confirm whether sequences identified from the three best represented taxonomic groups (*Nannochloropsis*, *Mycobacterium*, human) were likely to be of ancient origin or to have derived from modern contamination. The sequences aligned to the human genome did not exhibit typical patterns of ancient DNA damage, which include cytosine deamination and depurination-induced strand breaks and are therefore considered to be of modern contaminant origin (Supplementary Figure S2). In contrast, we find that sequences aligned to two *Nannochloropsis limnetica* and the *Mycobacterium avium* genomes exhibit authentic ancient DNA damage (Supplementary Figures S3 and S4), with patterns that are near identical for both taxonomic groups, consistent with their preservation in the same environment of broadly contemporaneous age.

We next aligned our sequence data against two organellar reference panels consisting of either 2742 chloroplast or 8486 mitochondrial genomes (Supplementary Table S5). Both analyses also recovered *N. limnetica* as the best represented taxon, with 23,600-37,900 and 8,600-14,100 sequences aligning to the chloroplast and mitochondrial genome of this taxon, respectively. After mapping the filtered sequence data to the *Nannochloropsis* chloroplast genomes individually, the number of sequences uniquely aligned to *N. limnetica* fell to 7,700-11,300 (Supplementary Table S4). The most abundant non-*Nannochloropsis* taxon was the algae *Choricystis parasitica* with 255-1,784 and 38-276 of sequences mapped to its chloroplast and mitochondrial genome.

### Reconstruction of *Nannochloropsis* organellar palaeogenomes and their phylogenetic placement

We reconstructed complete composite organellar palaeogenomes for *Nannochloropsis* present in LGM Andøya, using *N. limentica* as a seed sequence. The resulting complete chloroplast sequence was 117.7 kilobases (kb) in length and had a coverage depth of 64.3x. The mitochondrial genome was 38.5 kb in length with a coverage of 62.4x (Figure 2; Supplementary Figure S5; Supplementary Table S6). We observed two major structural changes in our reconstructed chloroplast as compared to the *N. limnetica* seed sequence, in which the reconstructed chloroplast was inferred to share the ancestral structural state with the remaining *Nannochloropsis* taxa. This included a 233 bp region in a non-coding region between the *thiG* and *rpl27* genes, which is absent in the *N. limnetica* seed sequence and of varying length among all other *Nannochloropsis* taxa (Supplementary Figure S6). A 323 bp insertion in a non-coding region between the genes *rbcS* and *psbA*, is present in the *N. limnetica* seed sequence, but lacking from our reconstructed chloroplast and all other *Nannochloropsis* taxa (Supplementary Figure S6). We noted that the combined coverage was reduced across these two regions, with 21x and 32x chloroplast genome coverage (the latter calculated from 100 bp upstream and downstream of the deletion), which may be suggestive of within-sample variation. Both the reconstructed composite organellar genomes displayed authentic ancient DNA damage patterns (Supplementary Figures S7 and S8).

**Figure 2:**
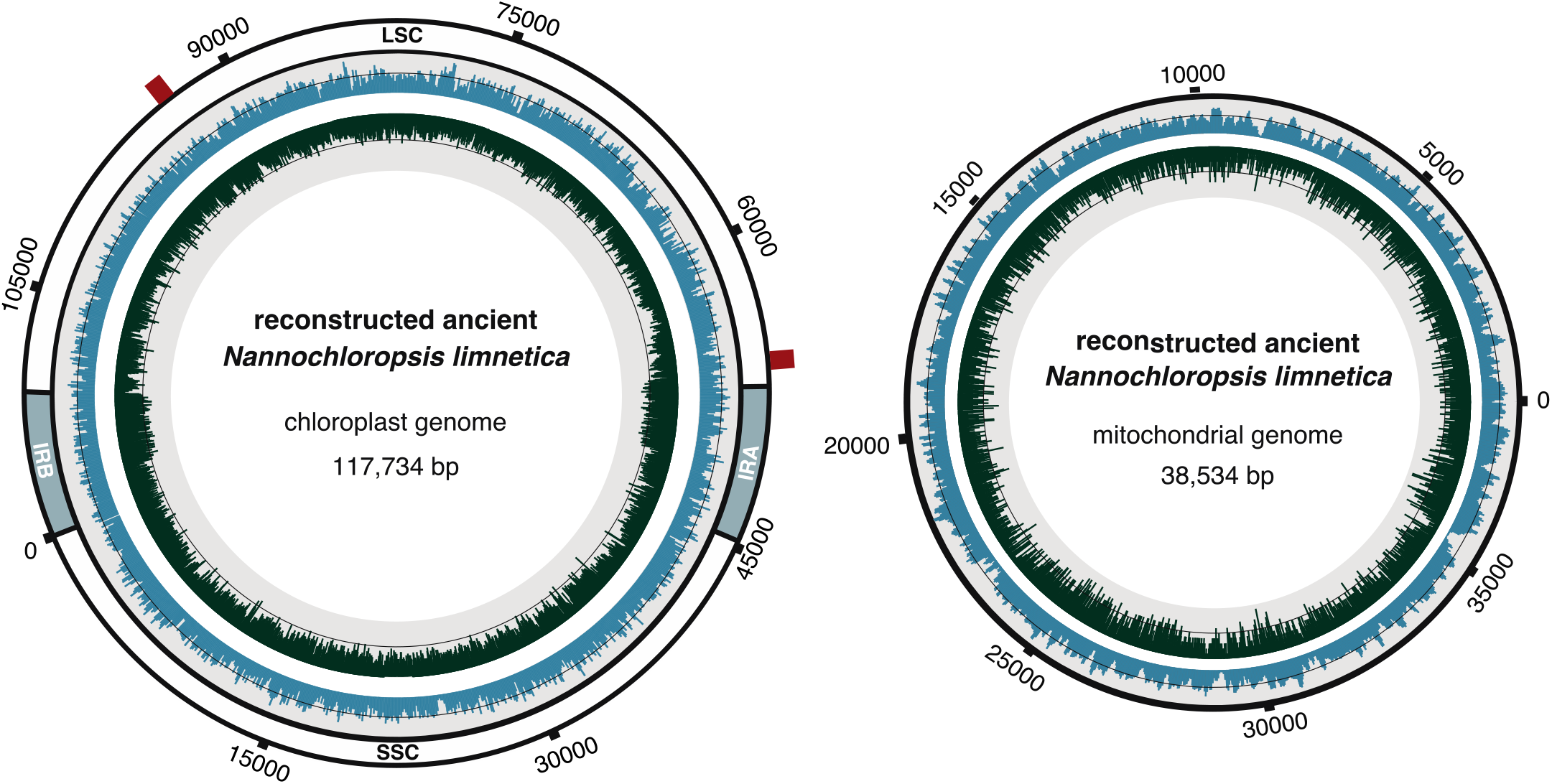
*N. limnetica* chloroplast and mitochondrial palaeogenomes reconstructed directly from *sed*aDNA. The innermost circle contains a distribution of the GC content in dark green, with the black line representing the 50% mark. The outer blue distribution contains the genomic coverage for the assembly, with the black line representing the average coverage of 64.3x for the chloroplast and 64.9x for the mitochondria. For the chloroplast the inverted repeats (IRA and IRB), large single copy (LSC) and small single copy (SSC) regions are annotated. The red bars on the chloroplast indicate the location of the two regions with structural change compared to the *N. limnetica* reference genome.

To account for within-sample variants in our reconstructed organellar palaeogenomes, we created two consensus sequences that included either high or low frequency variants at multiallelic sites. We performed phylogenetic analyses to confirm the placement of the high and low frequency variant consensus genomes relative to other *Nannochloropsis* taxa. For this, we used full organellar genomes and three short loci with high taxonomic representation in NCBI Genbank (18S, ITS, *rbc*L; Table S7). Altogether, these analyses from three different markers (chloroplast, mitochondrial, nuclear) were congruent and resolved the high frequency variant consensus sequences as likely deriving from *N. limnetica* var. *globosa* and the low frequency variant consensus sequences as *N. limnetica* var. *limnetica* (Table 2; Figure 3; Supplementary Figures S9).

**Figure 3:**
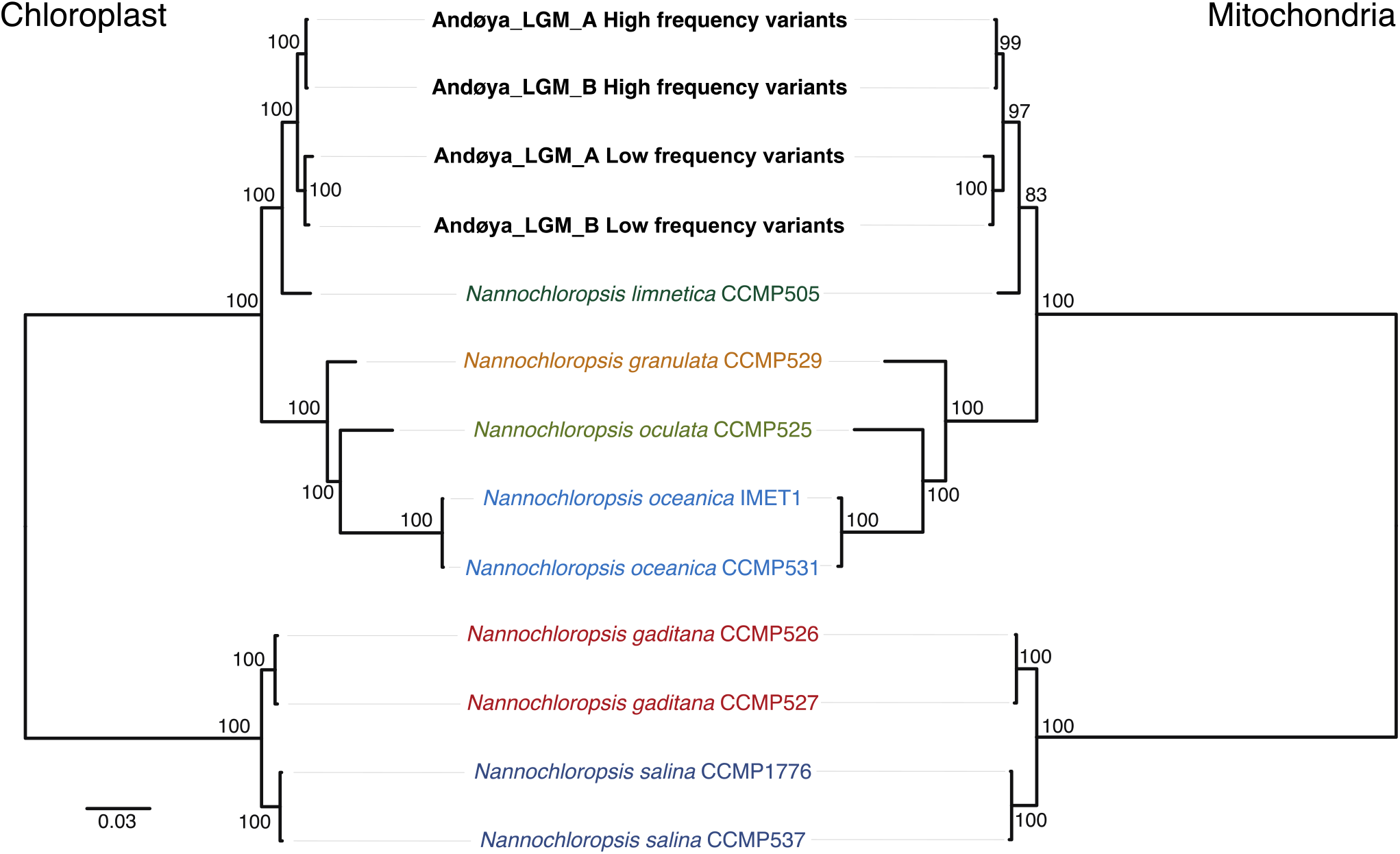
Maximum likelihood phylogeny of *Nannochloropsis* chloroplast (left) and mitochondrial (right) genome sequences, including the reconstructed *N. limnetica* consensus sequences based on either high or low frequency variants.

**Table 2:**
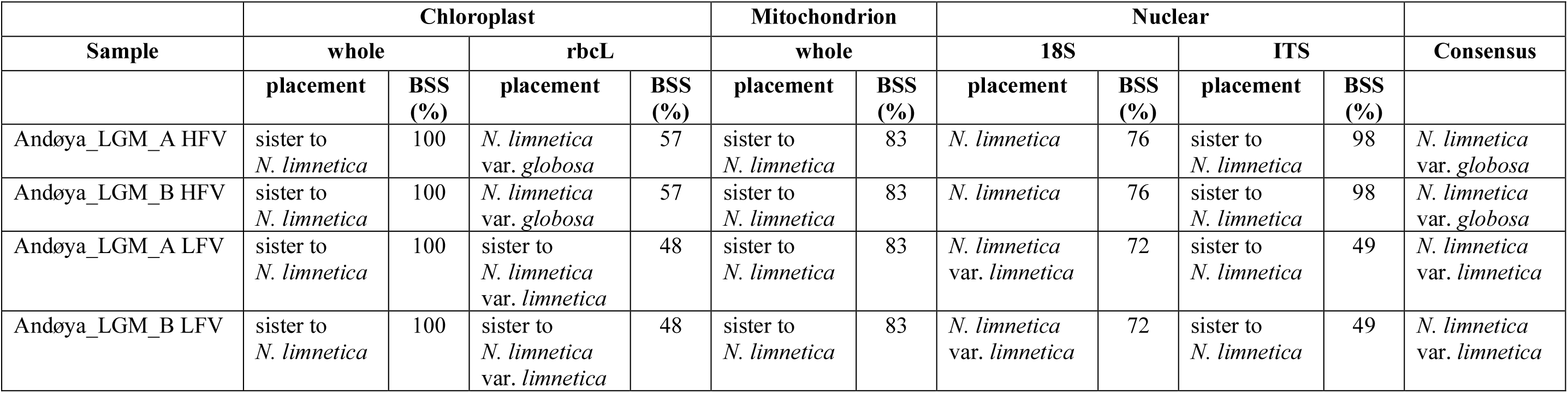
Placement of the reconstructed high and low frequency organellar genomes and markers in each phylogenetic analysis. BSS=Bootstrap support values, HFV=High frequency variant, LFV=Low frequency variant.

We attempted to reconstruct composite chloroplast genomes using alternative *Nannochloropsis* taxa as seed sequences, but these analyses failed to resolve a complete composite sequence (Supplementary Table S8). A phylogenetic analysis of these alternative composite chloroplast genomes displays a topology consistent with the biases associated with mapping to increasingly diverged reference genomes (Supplementary Figure S10). These alternative composite chloroplast genomes were therefore not used further, but provide supporting evidence that *N. limnetica* is the most closely related extant taxon.

### *Nannochloropsis limnetica* allelic variation and haplotype estimation

In the absence of a catalog of chloroplast and mitochondrial genomes from the *N. limnetica* variants, we sought to explore the frequencies and proportions of allelic variants present in our data set. We combined all sequences aligned to the high and low frequency variant consensus genomes into a single data set for each sample. We restricted our analyses to transversion variants only in order to exclude artifacts derived from ancient DNA damage, and defined the reference allele as that present in the reconstructed composite organellar genomes. We detected 299-376 and 81-112 variants within the *N. limnetica* chloroplast and mitochondrial genomes, respectively (Supplementary Table S9). For each sample and across the entire organellar genome, the average proportion of the transversion-only alternative allele is 0.39-0.42 for chloroplast variants and 0.39-0.43 for mitochondrial variants (Figure 4).

**Figure 4:**
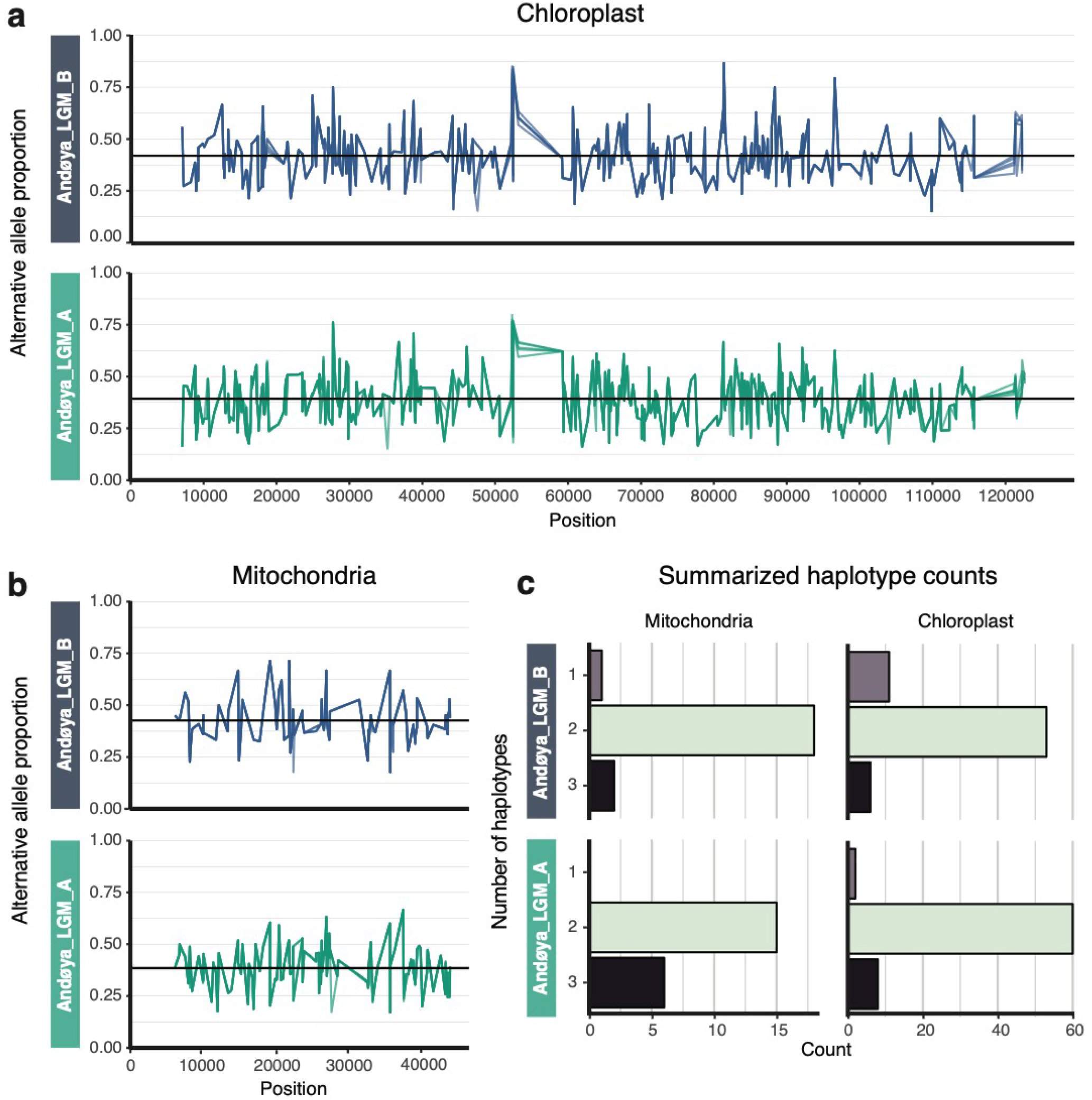
The proportions of alternative alleles across the chloroplast (a) and mitochondrial (b) genomes based on transversions-only. Each proportion plot consists of five independent variant calling runs to account for sampling biases (see Methods). The horizontal black lines represent averages: 0.39 and 0.42 for the chloroplast and 0.39 and 0.43 for the mitochondria, for samples Andøya_LGM_A and Andøya_LGM_B respectively. In (a) and (b), colour denotes sample. (c) Observed minimum haplotype counts based on the linked alleles for the chloroplast and mitochondrial genomes for both samples.

After pooling data from both samples, we used the phasing of adjacent alleles, which were linked by the same read, to infer the minimum number of haplotypes in each reconstructed composite organellar genome. We identify 70 and 21 transversion-only phased positions in the chloroplast and mitochondrial genomes, respectively. Within each sample, the average number of haplotypes observed, based on the linked alleles in the chloroplast genome, is 1.93-2.09. The equivalent average for the mitochondrial genome is 2.05-2.29 (Figure 4; Supplementary Table S10).

## Discussion

All of our analyses identified *Nannochloropsis* as the most abundant eukaryotic taxon in the LGM lake sediments from Andøya, consistent with a previous study based on plant DNA metabarcoding ^35^. We observed ancient DNA deamination patterns for all reference sequence combinations, which supports the authenticity of our data. The phylogenetic placement of our organellar palaeogenomes, as well as other short loci, indicate that the *Nannochloropsis* taxon detected in Andøya is *N. limnetica*, with at least two varieties present: *N. limnetica* var. *globosa* and *N. limnetica* var. *limnetica*.

The low overall proportion of sequences identified by the NCBI-based metagenomic analysis is broadly consistent with other shotgun metagenomic studies from *sed*aDNA ^10,13,14^ and suggests that the vast majority of taxonomic diversity in the sediment record is currently unidentifiable. We recovered comparable and low relative mapping frequencies for all non-*Nannochloropsis* eukaryotic taxa in our genome panel, regardless of their plausibility of occurring at LGM Andøya. We therefore suggest that these mappings are artifacts resulting from the spurious mapping of short and damaged ancient DNA molecules coupled with the vast diversity of sequences present in *sed*aDNA^53,54^. However, we identified a component of *Mycobacterium* sequences, which display ancient DNA damage patterns, although, unlike *Nannochloropsis*, no dominant strain could be identified. This indicates that our samples contain one or multiple unsequenced strains, some or all of which may be extinct.

We explored whether Andøya *Nannochloropsis* could potentially comprise more than one species. The detection of a low number of uniquely mapped sequences to non-*N. limnetica Nannochloropsis* genomes indicates that we cannot exclude the possibility of other rare *Nannochloropsis* taxa being present in the sequence data. However, we suggest that *N. limnetica* is the sole taxon present and that evolutionary divergence and potential technical artifacts can explain the conflicting results. Our phylogenetic analyses suggest that the Andøya *N. limnetica* variants are not evolutionarily close to any available organellar reference genomes. We therefore might expect some regions of our *N. limnetica* organellar genomes to be evolutionarily closer to non-*N. limnetica* than to *N. limnetica*. This is supported by the trend of decreasing quality of, and increased impact of reference bias upon, the chloroplast genome sequences reconstructed using alternative seed genomes. This could have been compounded by the aforementioned artifacts associated with ancient DNA ^53,54^.

We detected within-sample allelic variation in both the *N. limnetica* chloroplast and mitochondrial genome reconstructions, which we split into two consensus sequences containing either the high or the low frequency variants. In both organellar genomes, the high frequency variants for both samples, assigned as *N. limnetica* var. *globosa*, clustered separately from the low frequency variants, which we identified as *N. limnetica* var. *limnetica*. This demonstrates that the results from each of our broadly contemporaneous samples are replicable, which is further confirmed by comparable results obtained by the estimation of the minimum number of haplotypes. For both samples, our analyses recover at least two haplotypes, with a small proportion of phased positions containing three. We note that our method is conservative, given the strict filtering criteria and limited window size, and almost certainly underestimates true haplotype diversity. To accurately estimate the diversity and proportions of haplotypes, it is likely that an extensive reference database of *N. limnetica* haplotypes will be required. However, this may be particularly problematic for taxa that lack body fossils, which are currently required to reconstruct extinct haplotypes. Future methodological and statistical advances will therefore be required to estimate and quantify haplotype variation for taxa from within a *sed*aDNA population sample.

The sheer abundance of *N. limnetica* sequences in our identified *sed*aDNA shotgun sequence data suggests a high biomass of this algae in Lake Øvre Æråsvatnet during the LGM. *Nannochloropsis* is known to undergo blooming events that can reach up to 10^10^ cells per litre of water ^55^, which have been reported for *N. gaditana* in the Comacchio Lagoons, Italy ^56^ and *N. granulata* in the Yellow Sea, China ^55^. *N. limnetica* itself was first described from spring blooms in Germany, reaching concentrations up to 5.7×10^9^ cells per litre ^41^. Such blooms could explain the observed high sequence abundance in our data. Independent proxies from the same LGM sediments, including high loss-on-ignition (LOI) values ^32^ and organic elemental (C/N) proportions ^35^, are consistent with a blooming scenario resulting from high nutrient input. Stable isotope data suggest that the C/N is of a high trophic origin, most likely bird guano from an adjacent bird cliff ^35^, which corresponds with the detection of bird bones (little auk, *Alle alle*) in the LGM sediments ^30,33,35^. The high inflow of nutrients into the lake could have resulted in eutrophication of the lake ecosystem and thus initiated blooms of *N. limnetica*.

*Nannochloropsis* has not previously been reported from contemporary northern Norway, based on available Global Biodiversity Information Facility (GBIF) records and the published literature, which could be due to the general difficulty of observing and identifying this algae ^47,50^. We note, however, that our reanalysis of modern DNA metabarcoding data from 11 north Norwegian localities ^57^ shows the presence of *Nannochloropsis* at five sites (Supplementary Table S11), with dominant abundances detected for two sites. In addition to LGM Andøya ^35^, *Nannochloropsis* has either been previously reported, or unreported but present based on our re-analysis, in eight *sed*aDNA-based palaeoecological records from Greenland ^58^, St. Paul Island, Alaska, USA ^9,11^, Alberta, Canada ^10^, Latvia ^59^, Qinghai, China ^60^ and Svalbard ^52,61^(Supplementary Figure S11). We failed to detect *Nannochloropsis* in the Hässeldala Port, Sweden ^13^ record. We note that *Nannochloropsis* is particularly well represented in late Pleistocene and early Holocene sediments from these records, at a time when the climate was cooler than present ^62^. Assuming these records reflect *N. limnetica*, or an ecological analogue, then these occurrences are consistent with its known climatic tolerances, such as thriving in cold water ^43^. Therefore, climate would have been adequate for *N. limnetica* at Andøya during the LGM, whereas the high nutrient input could have stimulated the unusually high concentrations.

As *Nannochloropsis* taxa differ in their salinity tolerances, the ability to detect and identify them to the species-level could potentially be used as a palaeoecological proxy to estimate the salinity of coastal marine-lacustrine sedimentary records. The detection of the fresh or brackish water *N. limnetica* in Lake Øvre Æråsvatnet is consistent with earlier studies that indicated a lacustrine LGM sediment record ^25,32,34^. However, caution should be applied when assuming ecological preferences of a taxon that is evolutionarily divergent from reference sequences, as is the case here.

Our complete *N. limnetica* chloroplast palaeogenome reconstructions represent the first derived from *sed*aDNA, although a near-complete chloroplast sequence has recently been reported for a vascular plant ^14^. Although mitochondrial palaeogenomes have previously been reconstructed from cave sediments ^12^, and archaeological middens and latrines ^24,63^, ours are the first derived from lake sediments. The high depth of coverage for our sample-combined palaeogenomes (62-64x) allowed us to explore allelic proportions and haplotype diversity using *sed*aDNA, which resulted in us identifying at least two distinct variants. Together with recently published and ongoing studies, our work demonstrates the feasibility of the *sed*aDNA field moving into a new phase of environmental palaeogenomics. This will enable a broad range of ecological and evolutionary questions to be addressed using population genomic approaches, including for communities of taxa that may or may not be preserved in the body fossil record. With further innovations, this approach could also be extended to a suite of broad groups, including plants, invertebrates, and vertebrates, from lake catchments, cave sediments, and archaeological settings, therefore unlocking the full potential of *sed*aDNA.

## Material and methods

### Site description, chronology, and sampling

A detailed description of the site, coring methods, age-depth model reconstruction, and sampling strategy can be found in ^35^. Briefly, Lake Øvre Æråsvatnet is located on Andøya, Northern Norway (69.25579°N, 16.03517°E) (Figure 1). In 2013, two cores were collected from the deepest sediments, AND10 and AND11. Macrofossil remains were dated, with those from AND10 all dating to within the LGM. For the longer core AND11, a Bayesian age-depth model was required to estimate the age of each layer ^35^. In this study, we selected one sample of LGM sediments from each of the two cores. According to the Bayesian age-depth model, sample Andøya_LGM_B, from 1102 cm depth in AND11, was dated to a median age of 17,700 (range: 20,200-16,500) cal yr BP. The age of Andøya_LGM_A, from 938 cm depth in AND10, was estimated at 19,500 cal yr BP, based on the interpolated median date between two adjacent macrofossils (20 cm above: 19,940-18,980 cal yr BP, 30 cm below: 20,040-19,000 cal yr BP). As Andøya_LGM_A falls within the age range of Andøya_LGM_B, we consider the samples to be broadly contemporaneous.

### Sampling, DNA extraction, library preparation, and sequencing

The two cores were subsampled at the selected layers under clean conditions, in a dedicated ancient DNA laboratory at The Arctic University Museum of Norway in Tromsø. We extracted DNA from 15 g of sediment following the Taberlet phosphate extraction protocol ^18^ in the same laboratory. We shipped a 210 μL aliquot of each DNA extract to the ancient DNA dedicated laboratories at the Centre for GeoGenetics (University of Copenhagen, Denmark) for double-stranded DNA library construction. After concentrating the DNA extracts to 80 μL, half of each extract (40 μL, totalling between 31.7-36.0 ng of DNA) was converted into Illumina-compatible libraries using established protocols ^10^. The number of indexing PCR cycles was determined using qPCR and each sample was dual indexed. The libraries were then purified using the AmpureBead protocol (Beckman Coulter, Indianapolis, IN, USA), adjusting the volume ratio to 1:1.8 library:AmpureBeads, and quantified using a BioAnalyzer (Agilent, Santa Clara, CA, USA). The indexed libraries were pooled equimolarly and sequenced on a lane of the Illumina HiSeq 2500 platform using 2x 80 cycle paired-end chemistry.

### Raw read filtering

For each sample, we merged and adapter-trimmed the paired-end reads with *SeqPrep* (https://github.com/jstjohn/SeqPrep/releases, v1.2) using default parameters. We only retained the resulting merged sequences, which were then filtered with the preprocess function of the *SGA toolkit* v0.10.15 ^64^ by the removal of those shorter than 35 bp or with a DUST complexity score >1.

### Metagenomic analysis of the sequence data

We first sought to obtain an overview of the taxonomic composition of the samples and therefore carried out a BLAST-based metagenomic analysis on the two filtered sequence datasets. To make the datasets more computationally manageable, we subsampled the first and last one million sequences from the filtered dataset of each sample and analysed each separately. The data subsets were each identified against the NCBI nucleotide database (release 223) using the blastn function from the *NCBI-BLAST*+ suite v2.2.18+ ^65^ under default settings. For each sample, the results from the two subsets were checked for internal consistency, merged into one dataset, and loaded into *MEGAN* v6.12.3 ^66^. Analysis and visualization of the Last Common Ancestor (LCA) was carried out for the taxonomic profile using the following settings: min score=35, max expected=1.0E-5, min percent identity=95%, top percent=10%, min support percentage=0.01, LCA=naive, min percent sequence to cover=95%. We define sequences as the reads with BLAST hits assigned to taxa post-filtering, thus ignoring “unassigned” and “no hit” categories.

### Alignment to reference genome panels

We mapped our filtered data against three different reference panels to help improve taxonomic identifications and provide insight into the sequence abundance of the identified taxa (Supplementary Tables S4 and S5). The first reference panel consisted of 42 nuclear genomes that included taxa expected from Northern Norway, exotic/implausible taxa, human, six *Nannochloropsis* species, and four strains of *Mycobacterium*. The inclusion of exotic taxa was to give an indication of the background spurious mapping rate, which can result from mappings to conserved parts of the genome and/or short and damaged ancient DNA molecules ^53,54^. We included *Nannochloropsis*, *Mycobacterium*, and human genomes, due to their overrepresentation in the BLAST-based metagenomic analysis. The other two reference panels were based on either all mitochondrial or chloroplast genomes on NCBI GenBank (as of January 2018). The chloroplast data set was augmented with 247 partial or complete chloroplast genomes generated by the PhyloNorway project ^67^. The filtered data were mapped against each reference genome or organellar genome set individually using *bowtie2* v2.3.4.1^68^ under default settings. The resulting bam files were processed with *SAMtools* v0.1.19 ^69^. We removed unmapped sequences with *SAMtools view* and collapsed PCR duplicate sequences with *SAMtools rmdup*.

For the nuclear reference panel, we reduced potential spurious or nonspecific sequence mappings by comparing the mapped sequences to both the aligned reference genome and the NCBI nucleotide database using *NCBI-BLAST+*, following the method used by Graham *et al.* ^9^, as modified by Wang *et al.* ^11^. The sequences were aligned using the following *NCBI-BLAST*+ settings: num_alignments=100 and perc_identity=90. Sequences were retained if they had better alignments, based on bit score, to reference genomes as compared to the NCBI nucleotide database. If a sequence had a better or equal match against the NCBI nucleotide database, it was removed, unless the LCA of the highest NCBI nucleotide bit score was from the same genus as the reference genome (based on the NCBI taxonID). To standardize the relative mapping frequencies to genomes of different size, we calculated the number of retained mapped sequences per Mb of genome sequence.

The sequences mapped against the chloroplast and mitochondrial reference panels were filtered and reported in a different manner than the nuclear genomes. First, to exclude any non-eukaryotic sequences, we used *NCBI-BLAST*+ to search sequence taxonomies and retained sequences if the LCA was, or was within, Eukaryota. Second, for the sequences that were retained, the LCA was calculated and reported in order to summarize the mapping results across the organelle datasets. LCAs were chosen as the reference sets are composed of multiple genera.

Within the *Nannochloropsis* nuclear reference alignments, the relative mapping frequency was highest for *N. limnetica*. In addition, the relative mapping frequency for other *Nannochloropsis* taxa was higher than those observed for the exotic taxa. This could represent the mapping of sequences that are conserved between *Nannochloropsis* genomes or suggest the presence of multiple *Nannochloropsis* taxa in a community sample. We therefore cross-compared mapped sequences to determine the number of uniquely mapped sequences per reference genome. First, we individually remapped the filtered data to six available *Nannochloropsis* nuclear genomes, the accession codes of which are provided in Supplementary Table S4. For each sample, we then calculated the number of sequences that uniquely mapped to, or overlapped, between each *Nannochloropsis* genome. We repeated the above analysis with six available chloroplast sequences (Supplementary Table S4), to get a comparable overlap estimation for the chloroplast genome.

### Reconstruction of the Andøya *Nannochloropsis* community organellar palaeogenomes

To place the Andøya *Nannochloropsis* community taxon into a phylogenetic context, and provide suitable reference sequences for variant calling, we reconstructed environmental palaeogenomes for the *Nannochloropsis* mitochondria and chloroplast. First, the raw read data from both samples were combined into a single dataset and re-filtered with the *SGA toolkit* to remove sequences shorter than 35 bp, but retain low complexity sequences to assist in the reconstruction of low complexity regions in the organellar genomes. This re-filtered sequence data set was used throughout the various steps for environmental palaeogenome reconstruction.

The re-filtered sequence data were mapped onto the *N. limnetica* reference chloroplast genome (NCBI GenBank accession: NC_022262.1) with *bowtie2* using default settings. *SAMtools* was used to remove unmapped sequences and PCR duplicates, as above. We generated an initial consensus genome from the resulting bam file with *BCFtools* v1.9 ^69^, using the *mpileup*, *call*, *filter*, and *consensus* functions. For variable sites, we produced a majority-rule consensus using the --variants-only and --multiallelic-caller options, and for uncovered sites the reference genome base was called. The above steps were repeated until the consensus could no longer be improved. The re-filtered sequence data was then re-mapped onto the initial consensus genome sequence with *bowtie2*, using the above settings. The *genomecov* function from *BEDtools* v2.17.0 ^70^ was used to identify gaps and low coverage regions in the resulting alignment.

We attempted to fill the identified gaps, which likely consisted of diverged or difficult-to-assemble regions. For this, we assembled the re-filtered sequence dataset into *de novo* contigs with the MEGAHIT pipeline v1.1.4 ^71^, using a minimum *k*-mer length of 21, a maximum *k*-mer length of 63, and *k*-mer length increments of six. The MEGAHIT contigs were then mapped onto the initial consensus genome sequence with the *blastn* tool from the *NCBI-BLAST+* toolkit. Contigs that covered the gaps identified by *BEDtools* were incorporated into the initial consensus genome sequence, unless a *blast* comparison against the NCBI nucleotide database suggested a closer match to non-*Nannochloropsis* taxa. We repeated the *bowtie2* gap-filling steps iteratively, using the previous consensus sequence as reference, until a gap-free consensus was obtained. The re-filtered sequence data were again mapped, the resulting final assembly was visually inspected, and the consensus was corrected where necessary. This was to ensure the fidelity of the consensus sequence, which incorporated *de novo*-assembled contigs that could potentially be problematic, due to the fragmented nature and deaminated sites of ancient DNA impeding accurate assembly ^72^.

Annotation of the chloroplast genome was carried out with *GeSeq* ^73^, using the available annotated *Nannochloropsis* chloroplast genomes (accession codes provided in Supplementary Table S12). The resulting annotated chloroplast was visualised with *OGDRAW* ^74^.

The same assembly and annotation methods outlined above were used to reconstruct the mitochondrial palaeogenome sequence, where the initial mapping assembly was based on the *N. limnetica* mitochondrial sequence (NCBI GenBank accession: NC_022256.1). The final annotation was carried out by comparison against all available annotated *Nannochloropsis* mitochondrial genomes (accession codes provided in Supplementary Table S12).

If the *Nannochloropsis* sequences derived from more than one taxon, then alignment to the *N. limnetica* chloroplast genome could introduce reference bias, which would underestimate the diversity of the *Nannochloropsis* sequences present. We therefore reconstructed *Nannochloropsis* chloroplast genomes, but using the six available *Nannochloropsis* chloroplast genome sequences, including *N. limnetica*, as seed genomes (accession codes for the reference genomes are provided in Supplementary Table S8). The assembly of the consensus sequences followed the same method outlined above, but with two modifications to account for the mapping rate being too low for complete genome reconstruction based on alignment to the non-*N. limnetica* reference sequences. First, consensus sequences were called with *SAMtools*, which does not incorporate reference bases into the consensus at uncovered sites. Second, neither additional gap filling, nor manual curation was implemented.

### Assembly of high and low frequency variant consensus sequences

The within-sample variants in each reconstructed organellar palaeogenome was explored by creating two consensus sequences, which included either high or low frequency variants at multiallelic sites. For each sample, the initial filtered sequence data were mapped onto the reconstructed *Nannochloropsis* chloroplast palaeogenome sequence with *bowtie2* using default settings. Unmapped and duplicate sequences were removed with *SAMtools*, as above. We used the *BCFtools mpileup*, *call*, and *normalize* functions to identify the variant sites in the mapped dataset, using the --skip-indels, --variants-only, and --multiallelic-caller options. The resulting alleles were divided into two sets, based on either high or low frequency variants. High frequency variants were defined as those present in the reconstructed reference genome sequence. Both sets were further filtered to only include sites with a quality score of 30 or higher and a coverage of at least half the average coverage of the mapping assembly (minimum coverage: Andøya_LGM_A=22x, Andøya_LGM_B=14x). We then generated the high and low frequency variant consensus sequences using the consensus function in *BCFTools*. The above method was repeated for the reconstructed *Nannochloropsis* mitochondrial genome sequence in order to generate comparable consensus sequences of high and low frequency variants (minimum coverage: Andøya_LGM_A=16x, Andøya_LGM_B=10x).

### Analysis of ancient DNA damage patterns

We checked for the presence of characteristic ancient DNA damage patterns for sequences aligned to four nuclear genomes: human, *Nannochloropsis limnetica* and *Mycobacterium avium*. We further analysed damage patterns for sequences aligned to both the reconstructed *N. limnetica* composite organellar genomes. Damage analysis was conducted with *mapDamage* v2.0.8 ^75^ using the following settings: --merge-reference-sequences and --length=160.

### Phylogenetic analysis of the reconstructed organellar palaeogenomes

We determined the phylogenetic placement of our high and low frequency variant organellar palaeogenomes within *Nannochloropsis*, using either full mitochondrial and chloroplast genome sequences or three short loci (18S, ITS, *rbc*L). We reconstructed the 18S and ITS1-5.8S-ITS2 complex using DQ977726.1 (full length) and EU165325.1 (positions 147:1006, corresponding to the ITS complex) as seed sequences following the same approach that was used for the organellar palaeogenome reconstructions, except that the first and last 10 bp were trimmed to account for the lower coverage due to sequence tiling. We then called high and low variant consensus sequences as described above.

We created six alignments using available sequence data from NCBI Genbank (Supplementary Tables S7) with the addition of: (1+2) the high and low frequency variant chloroplast or mitochondrial genome consensus sequences, (3) a ~1100 bp subset of the chloroplast genome for the *rbc*L alignment, (4+5) ~1800 bp and ~860 bp subsets of the nuclear multicopy complex for the 18S and ITS alignments, respectively, and (6) the reconstructed chloroplast genome consensus sequences derived from the alternative *Nannochloropsis* genome starting points. Full details on the coordinates of the subsets are provided in Supplementary Table S7. We generated alignments using *MAFFT* v7.427 ^76^ with the maxiterate=1000 setting, which was used for the construction of a maximum likelihood tree in *RAxML* v8.1.12 ^77^ using the GTRGAMMA model and without outgroup specified. We assessed branch support using 1000 replicates of rapid bootstrapping.

### *Nannochloropsis* variant proportions and haplogroup diversity estimation

To estimate major haplogroup diversity, we calculated the proportions of high and low variants in the sequences aligned to our reconstructed *Nannochloropsis* mitochondrial and chloroplast genomes. For each sample, we first mapped the initial filtered sequence data onto the high and low frequency variant consensus sequences with *bowtie2*. To avoid potential reference biases, and for each organellar genome, the sequence data were mapped separately against both frequency consensus sequences. The resulting bam files were then merged with *SAMtools* merge. We removed exact sequence duplicates, which may have been mapped to different coordinates, from the merged bam file by randomly retaining one copy. This step was replicated five times to examine its impact on the estimated variant proportions. After filtering, remaining duplicate sequences - those with identical mapping coordinates - were removed with *SAMtools* rmdup. We then called variable sites from the duplicate-removed bam files using *BCFTools* under the same settings as used in the assembly of the high and low frequency variant consensus sequences. We restricted our analyses to transversion-only variable positions, to remove the impact of ancient DNA deamination artifacts. For each variable site, the proportion of reference and alternative alleles was calculated, based on comparison to the composite *N. limnetica* reconstructed organellar palaeogenomes. We removed rare alleles occurring at a proportion of <0.1, as these may have resulted from noise.

To infer the minimum number of haplogroups in each reconstructed organellar genome sequence, we inspected the phasing of adjacent variable sites that were linked by the same read in the duplicate-removed bam files, akin to the method used by Søe *et al*. ^63^. For this, we first identified all positions, from both samples, where two or more transversion-only variable sites occurred within 35 bp windows. We then examined the allelic state in mapped sequences that fully covered each of these linked positions. We recorded the combination of alleles to calculate the observed haplotype diversity at each of the linked positions. We removed low frequency haplotypes, which were defined as those with <3 sequences or <15% of all sequences that covered a linked position, and the remaining haplotypes were scored.

### Meta-analysis of *Nannochloropsis* in previous *sed*aDNA data sets

We performed a meta-analysis of the global prevalence of *Nannochloropsis* since the last ice age using published and available lake *sed*aDNA data sets. Three published shotgun datasets from Lake Hill, Alaska, USA ^9,11^, Charlie Lake, BC and Spring Lake, Alberta, Canada ^10^, and Hässeldala Port, Sweden ^13^ were reanalysed for the presence of *N. limnetica* using the same nuclear genome method as used in this study (Supplementary Table S13). Furthermore, a metabarcode data set was reanalysed from Skartjørna, Svalbard ^61^, using the same methods for analysis as the original study, but lowering the minimum barcode length to 10 bp, in order to retain the *Nannochloropsis* barcode (tag-sample lookup is provided in Supplementary Table S14). These data sets were supplemented with *sed*aDNA metabarcoding studies that reported *Nannochloropsis*, including; Bliss Lake, Greenland ^58^, Qinghai Lake, China ^60^, Lielais Svẽtiņu, Latvia ^59^, Lake Øvre Æråsvatnet ^35^, and Jodavannet, Svalbard ^52^.

We estimated the occurrence and abundance of *Nannochloropsis* in 5,000-year time windows for the above data sets. Abundance was coarsely divided into four categories for the metabarcode data: (1) dominant, scored when *Nannochloropsis* was the only taxon detected or most abundant of the taxa identified in the sequence data; (2) common, assigned when it was in the top 10 most abundant taxa identified; (3) rare, scored for any other detections, and (4) absent, assigned if *Nannochloropsis* was not detected. The reanalysed shotgun data sets were scored as: (1) dominant, when *Nannochloropsis* made up >=0.1% of the filtered read data; (2) common, 0.09-0.01%; (3) rare, 0.009-0.001%, and (4) absent, with <0.001%.

## Data availability

The raw Illumina shotgun sequence datasets are available from EMBL via *ACCESSION CODES*. The reanalysed metabarcoding data from Alsos et al. ^61^ are available via *ACCESSION CODE*. The reconstructed *Nannochloropsis limnetica* high and low frequency organellar genome sequences are available from NCBI Genbank via *ACCESSION CODES*. The scripts estimating the number of haplotypes across the linked windows are provided in the following GitHub repository at *GITHUB LINK*.

## Acknowledgements

This paper is a part of a larger project on the past environment of Andøya and we thank the Andøya team for fruitful discussion and access to pre-publication results. We thank Per Sjögren, Aage Paus and Ludovic Gielly for assistance with fieldwork; Antony G. Brown for help with the age-depth models; Mikkel W. Pedersen for conducting the library preparation and overseeing the sequencing; Edana Lord, Vendela K. Lagerholm, and Love Dalén for access to the pre-published *Lemmus lemmus* genome; and Sandra Garcés Pastor for informative discussions. The work was funded by the Research Council of Norway (grants: 213692, Ancient DNA of NW Europe reveals responses of climate change; 250963, ECOGEN – Ecosystems change and species persistence over time: a genome-based approach to IGA). YL was financed by an internal PhD position at The Arctic University Museum of Norway.

## Author contribution

YL, PDH, and IGA conceptualised and designed the research, and contributed to the final version of the manuscript; YL analysed the data and wrote the first draft; PDH provided analytical guidance and refined the drafted manuscript; IGA performed fieldwork, DNA extractions, provided resources, acquired funding, and supervised YL.

