## Supplementary Figures and Tables for "Environmental palaeogenomic reconstruction of an Ice Age algal population"

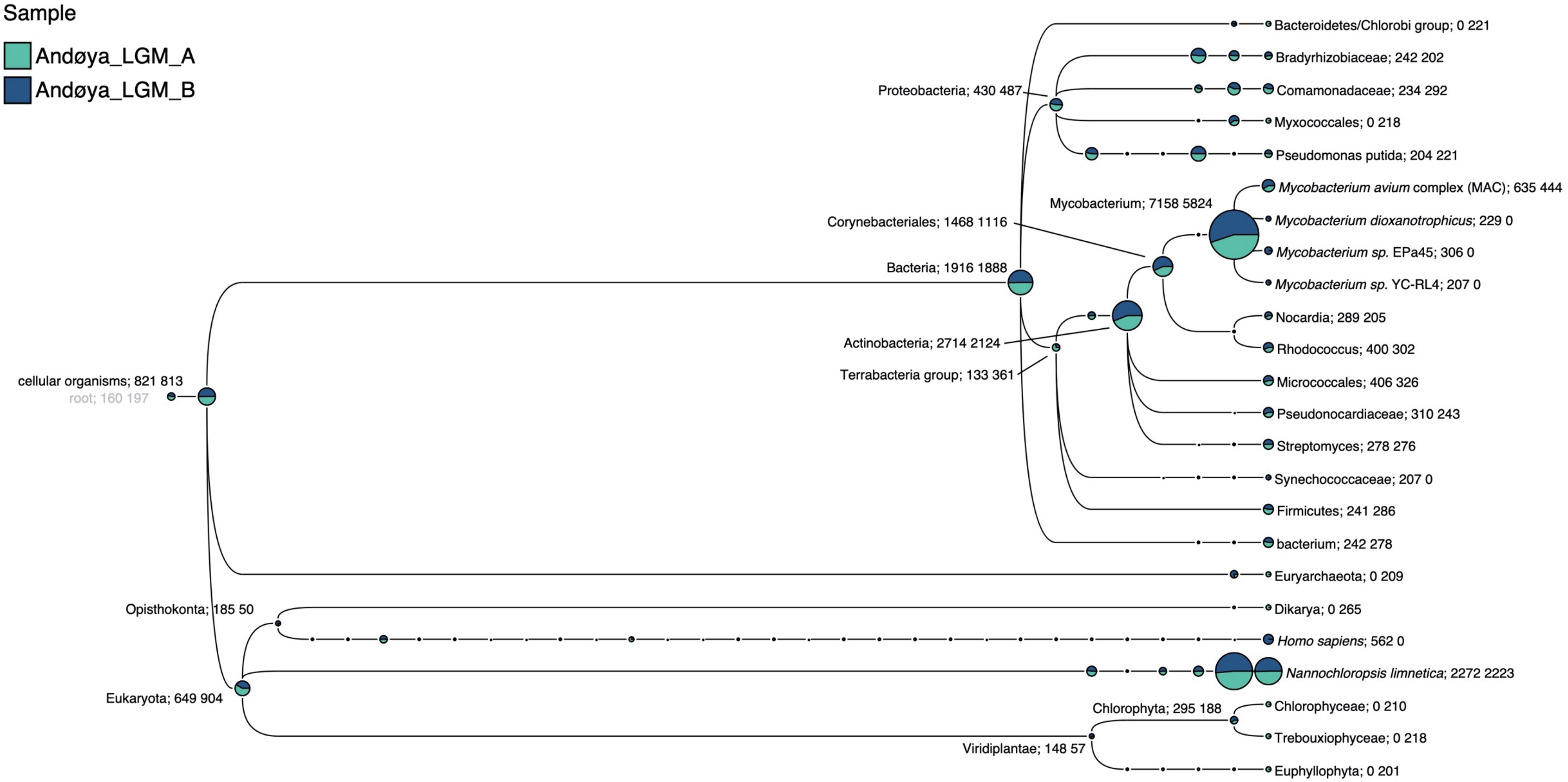


**Supplementary Figure S1**: Visualization of results from the BLAST-based metagenomic analysis. All tips are collapsed to the genus-level or higher, with the exception of the two most read-abundant tips: *Mycobacterium* and *Nannochloropsis*.

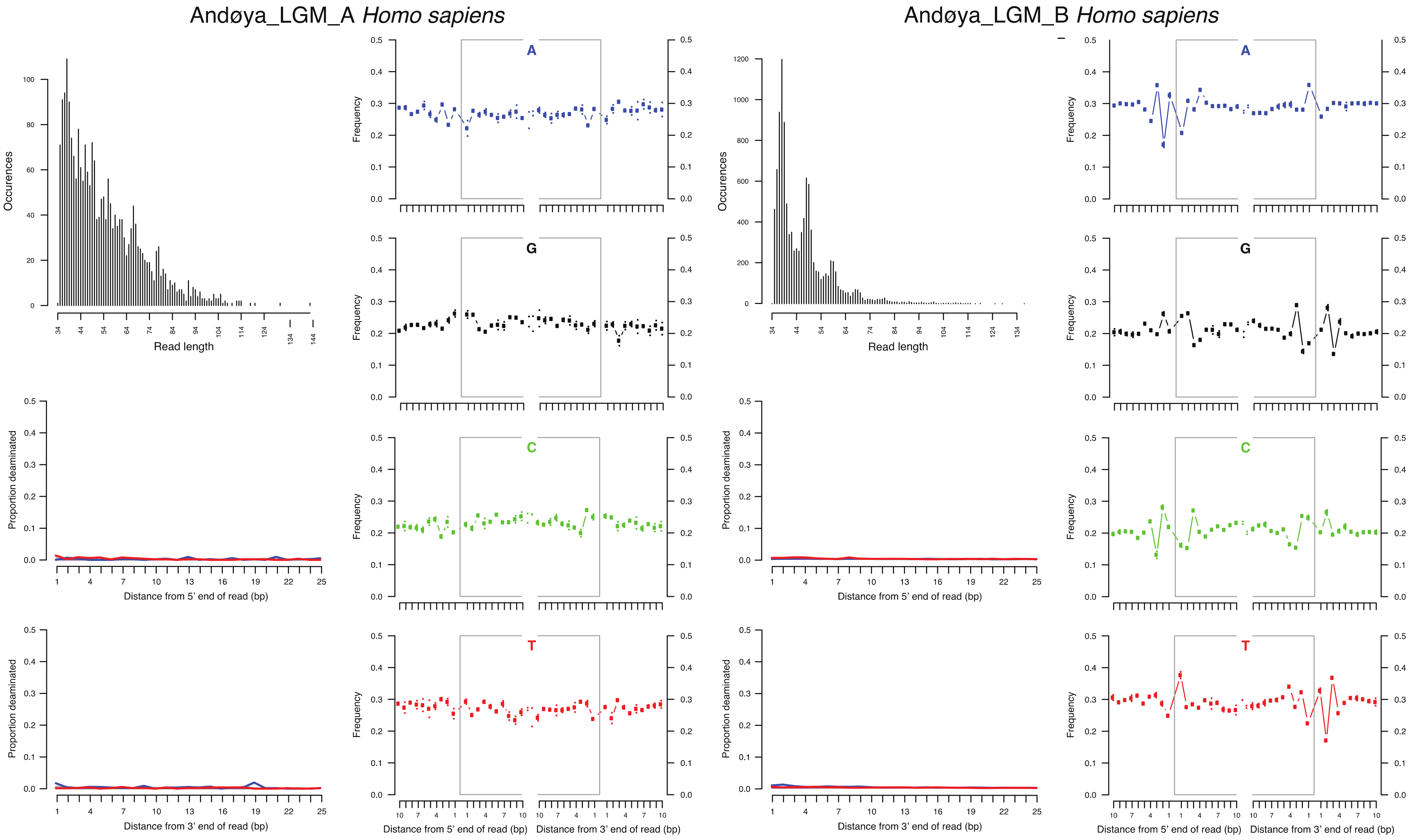


**Supplementary Figure S2**: mapDamage results for sequences aligned to the *Homo sapiens* nuclear reference genome.

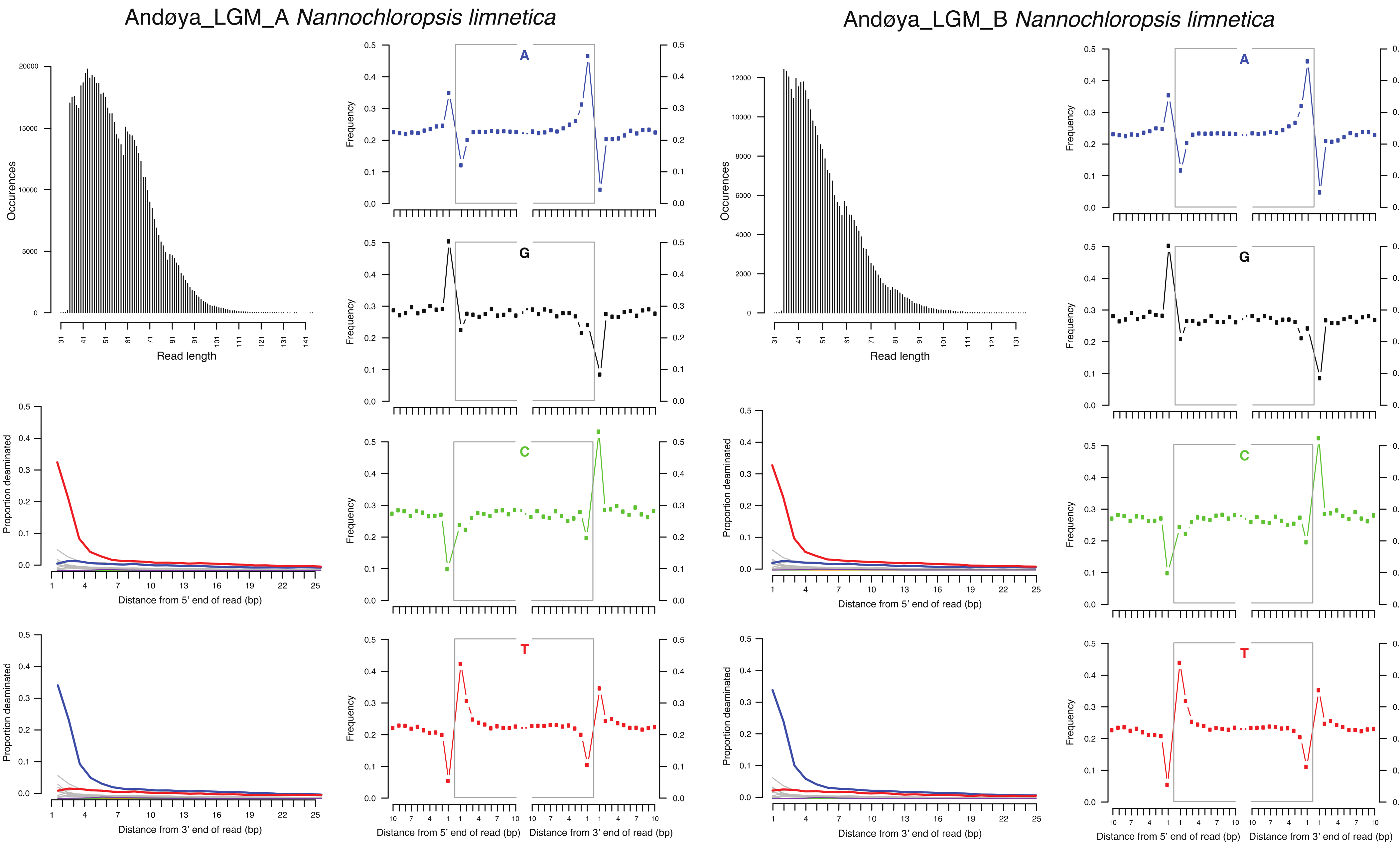


**Supplementary Figure S3**: mapDamage results for sequences aligned to the *Nannochloropsis limnetica* nuclear reference genome.

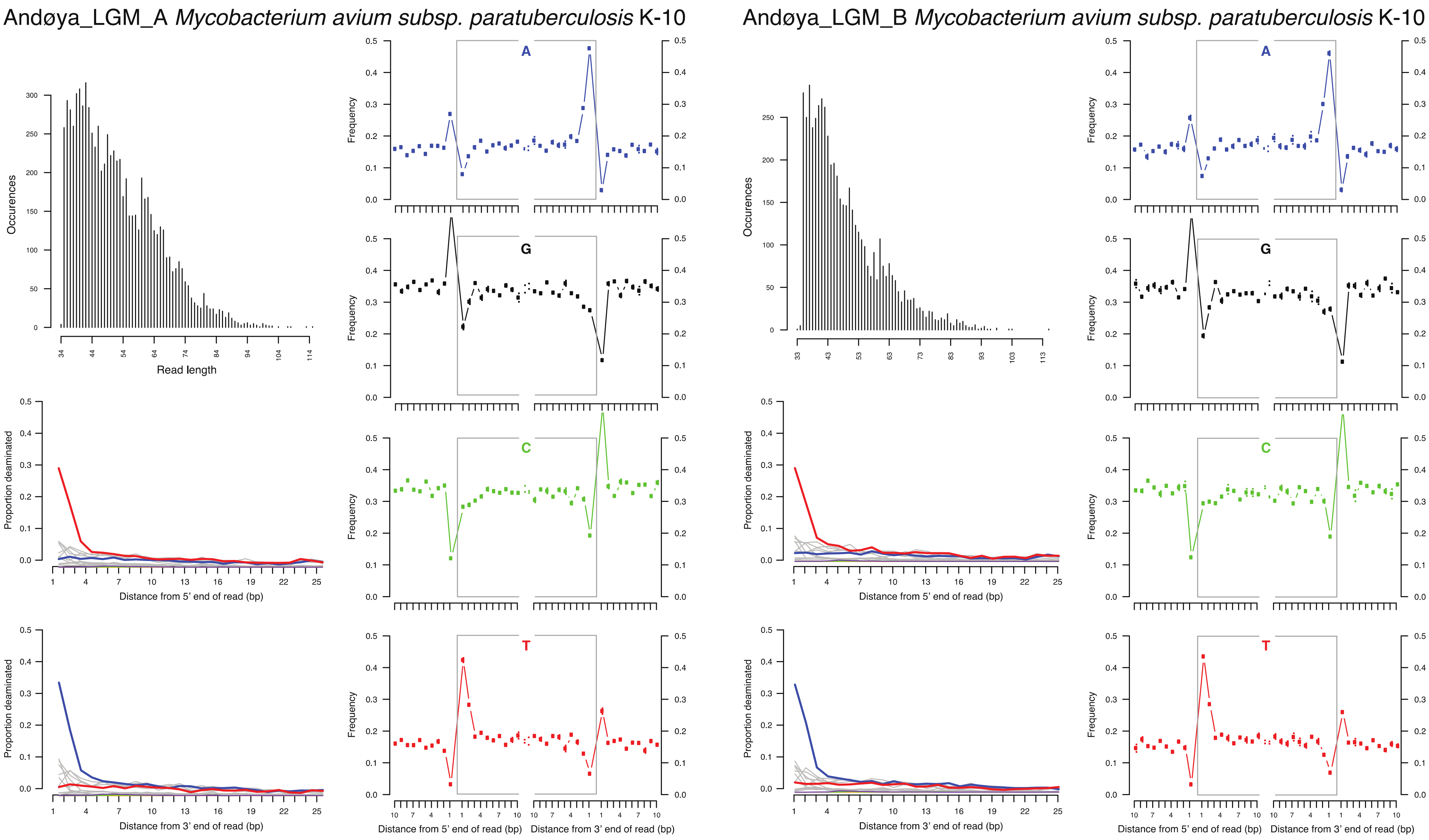


12 **Supplementary Figure S4**: mapDamage results for sequences aligned to the *Mycobacterium avium* nuclear reference genome.

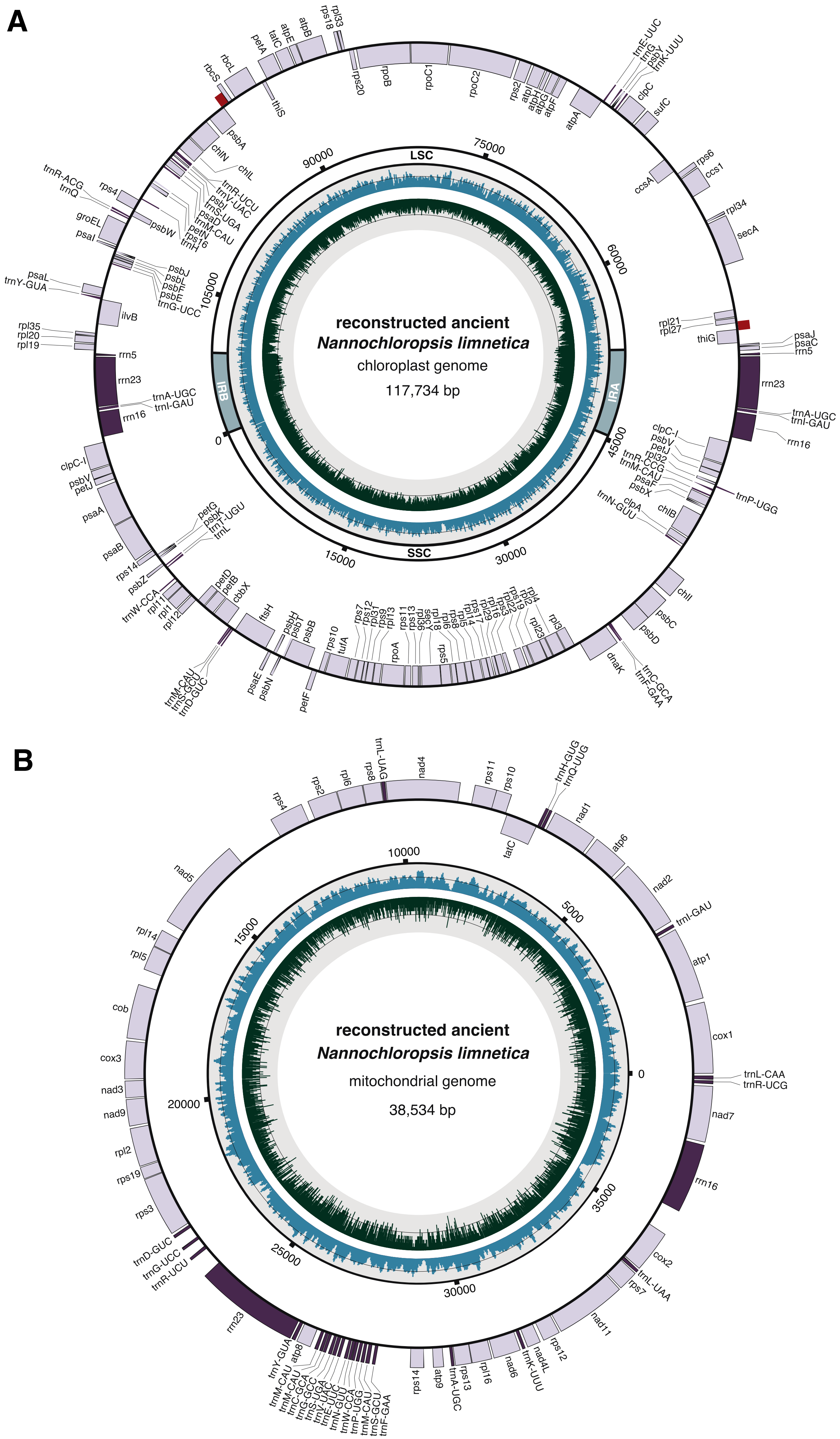

A
reconstructed ancient
Nannochloropsis limnetica
chloroplast genome
117,734 bp
LSC
SSC
IRA
IRB
0
15000
30000
45000
60000
75000
90000
105000
rbcS
rbcL
thiS
petA
tatC
atpE
atpB
rps18
rpl33
rps20
rpoB
rpoC1
rpoC2
rps2
atpI
atpH
atpG
atpF
atpA
trnE-UUC
trnG
psbY
trnK-UUU
clpC
sufC
rps6
ccs1
ccsA
rpl34
secA
rpl21
rpl27
thiG
psaJ
psaC
rrn5
rrn23
trnA-UGC
trnI-GAU
rrn16
clpC-I
psbV
petJ
rpl32
trnR-CCG
trnM-CAU
psaF
psbX
trnP-UGG
clpA
trnN-GUU
chlB
chlI
psbC
psbD
dnaK
trnC-GCA
trnF-GAA
psbA
chlN
chlL
trnR-UCU
trnV-UAC
psbI
trnS-UGA
psaD
petN
trnM-CAU
rps16
trnH
trnR-ACG
trnQ
rps4
psbW
groEL
psaI
psbJ
psbL
psbF
psbE
trnG-UCC
psaL
trnY-GUA
ilvB
rpl35
rpl20
rpl19
psaA
psaB
petG
psbK
trnT-UGU
trnL
rps14
psbZ
trnW-CCA
rpl11
rpl1
rpl12
petD
petB
cbbX
trnM-CAU
trnS-GCU
trnD-GUC
ftsH
psaE
psbN
psbH
psbT
psbB
petF
rps10
tufA
rps7
rps12
rpl31
rps9
rpl13
rpoA
rps11
rps13
rpl36
secY
rpl18
rps5
rpl6
rps8
rpl5
rpl14
rps17
rpl29
rpl16
rps3
rps19
rpl2
rpl4
rpl23
rpl3
B
reconstructed ancient
Nannochloropsis limnetica
mitochondrial genome
38,534 bp
0
5000
10000
15000
20000
25000
30000
35000
rps4
rps2
rpl6
rps8
trnL-UAG
nad4
rps11
rps10
tatC
trnH-GUG
trnQ-UUG
nad1
atp6
nad2
trnI-GAU
atp1
cox1
trnL-CAA
trnR-UCG
nad7
rrn16
cox2
trnL-UAA
rps7
nad11
rps12
nad4L
trnK-UUU
nad6
rpl16
rps13
trnA-UGC
atp9
rps14
trnF-GAA
trnS-GCU
trnM-CAU
trnP-UGG
trnW-CCA
trnN-GUU
trnE-UUC
trnV-UAC
trnS-UGA
trnG-GCC
trnC-GCA
trnM-CAU
trnY-GUA
atp8
rrn23
trnR-UCU
trnG-UCC
trnD-GUC
rps3
rps19
rpl2
nad9
nad3
cox3
cob
rpl5
rpl14
nad5

**Supplementary Figure S5**: Fully annotated maps for the reconstructed *N limnetica* (A) chloroplast and (B) mitochondrial palaeogenomes. The innermost circle contains a distribution of the GC content in dark green, with the black line representing the 50% mark. The outer distribution contains the coverage for the assembly in blue, with the black line representing the average coverage of 64.3x for the chloroplast and 64.9x for the mitochondria. For the chloroplast the inverted repeats (IRA and IRB), large single copy (LSC) and small single copy (SSC) regions are annotated. The genomic features are given on the outermost circle, where the coding genes are coloured light purple and RNAs in dark purple. The features located on the inside are transcribed clockwise, those on the outside anticlockwise. The red bars on the outermost circle of the chloroplast indicate the location of the two regions with structural change compared to the *N. limnetica* reference genome.

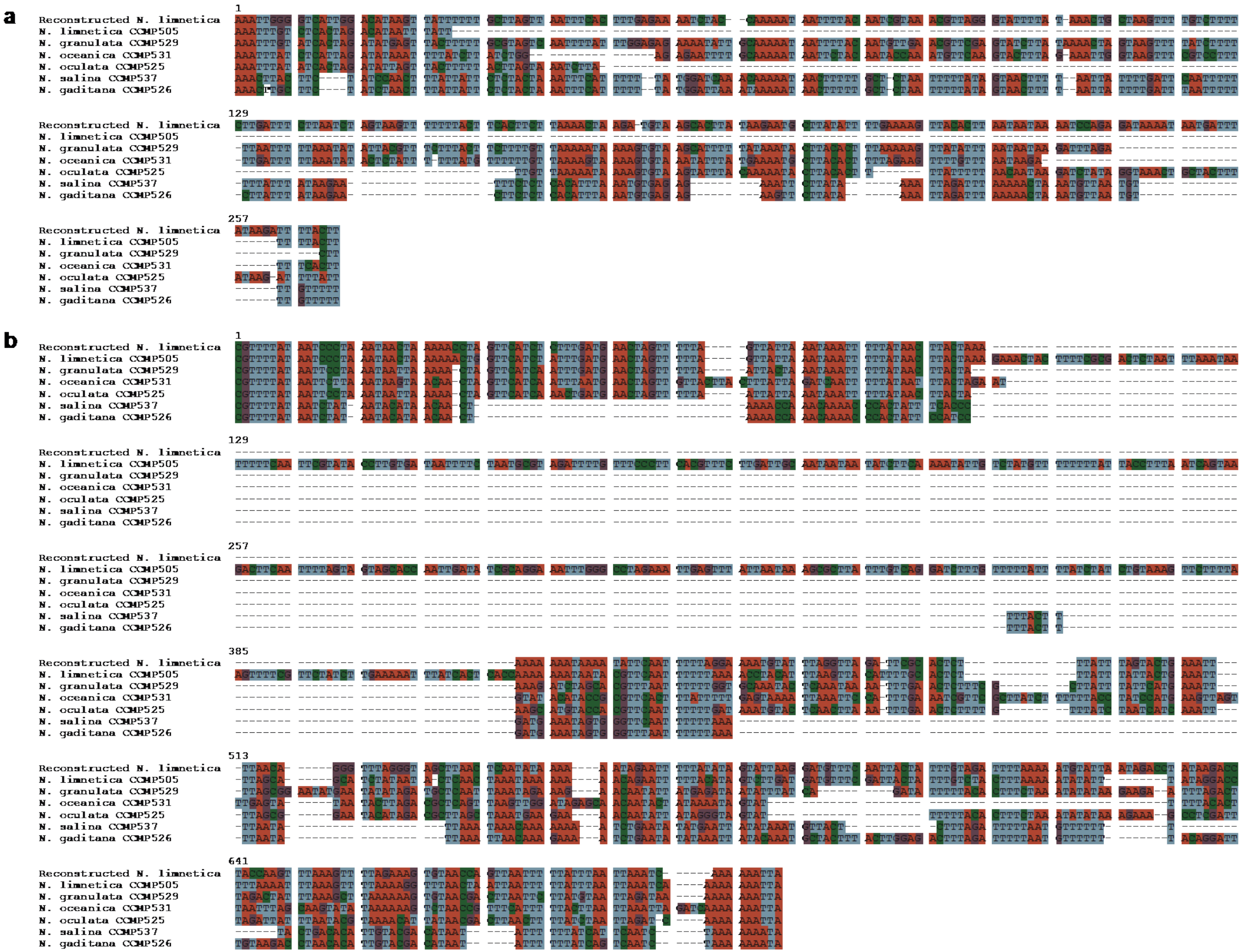


**Supplementary Figure S6**: Alignments for the two major structural changes in the reconstructed chloroplast compared to the six reference *Nannochloropsis* chloroplast genomes. (a) A 233 bp insertion in a non-coding region between the *thiG* and *rpl27* genes, which is absent in the reference *N. limnetica* chloroplast. (b) A 323 bp deletion compared to the reference *N. limnetica* chloroplast in a non-coding region between the *rbcS* and *psbA* genes. The accession codes for the *Nannochloropsis* sequences are provided in Supplementary Table S4.

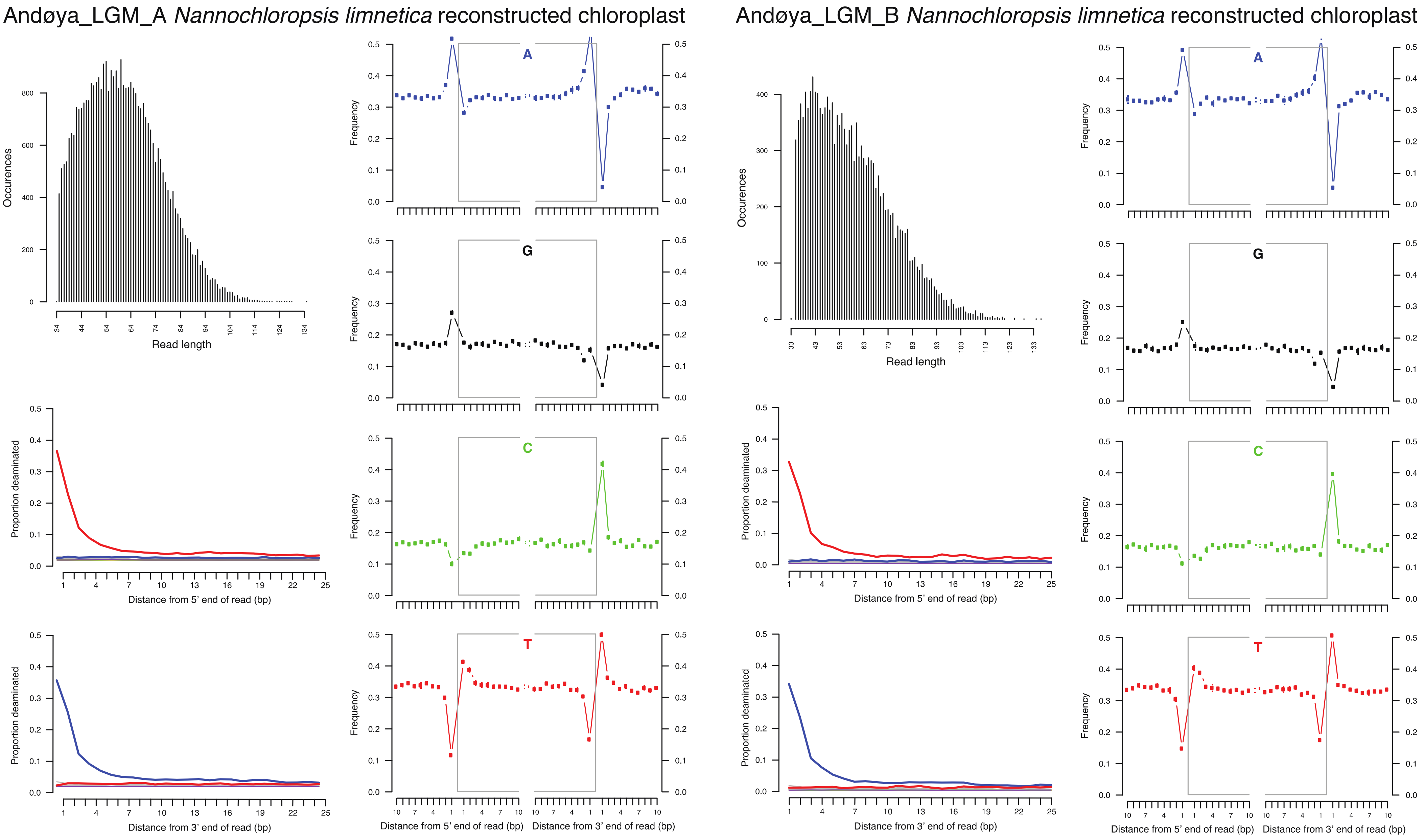


**Supplementary Figure S7**: mapDamage results for sequences aligned to the reconstructed *N. limnetica* chloroplast genome.

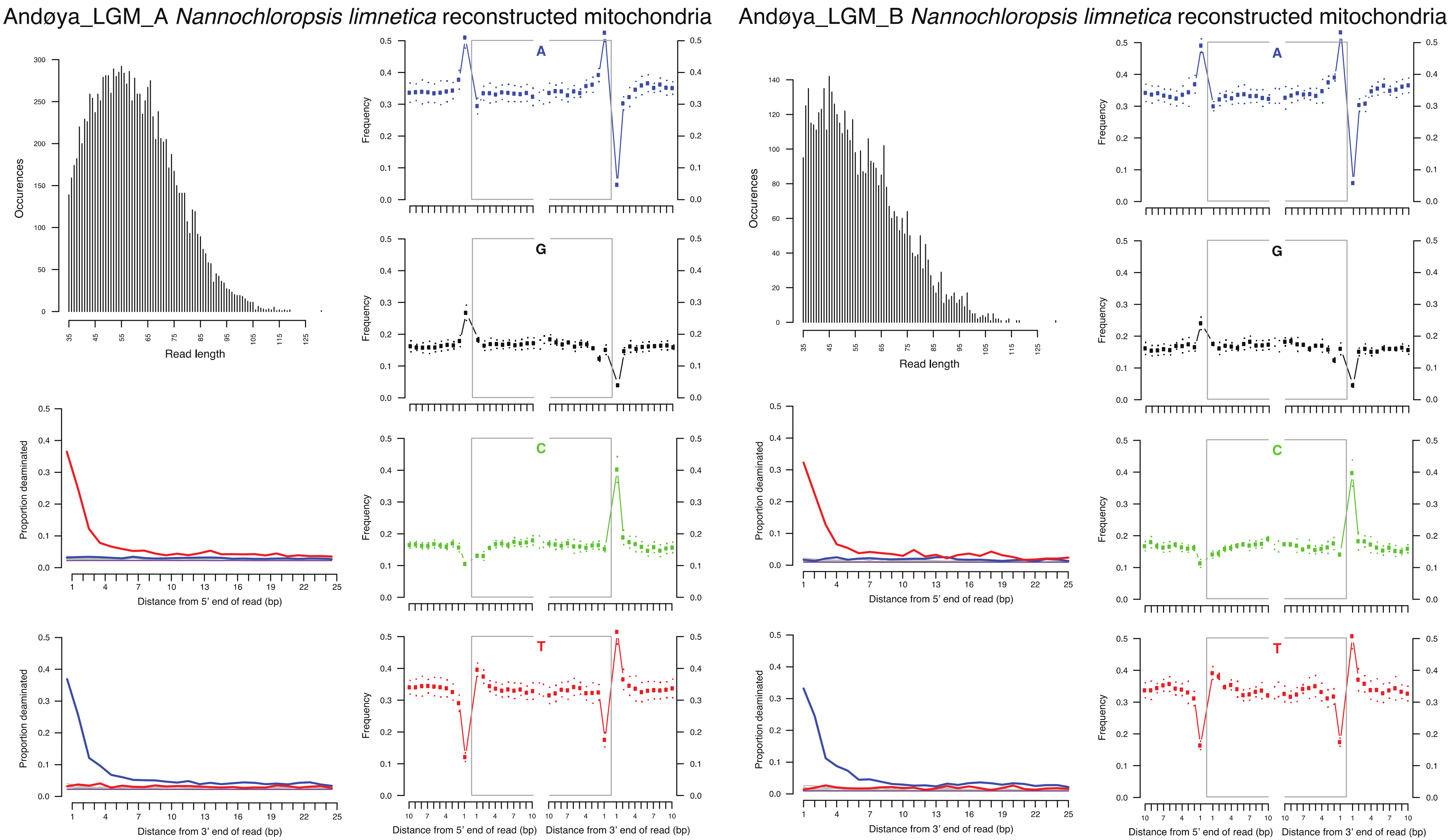


30 **Supplementary Figure S8**: mapDamage results for sequences aligned to the reconstructed *N. limnetica* mitochondrial genome.

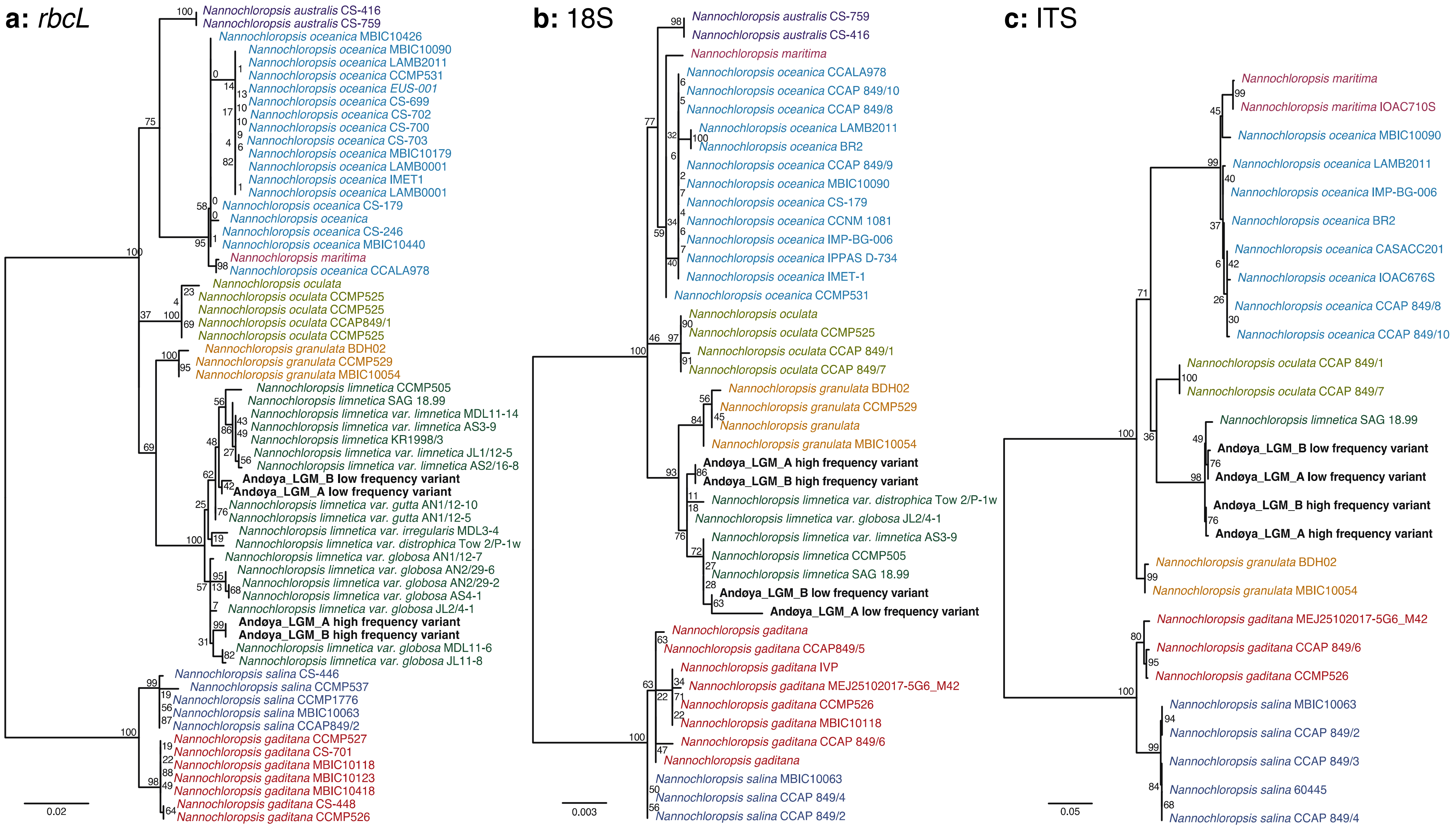


**Supplementary Figure S9**: Maximum likelihood phylogeny of *Nannochloropsis*, including the reconstructed *N. limnetica* high and low frequency variant consensus sequences, based on (a) ~1100 bp of the *rbcL* chloroplast locus, (b) ~1800 bp of the 18S nuclear locus, and (c) ~860 bp of the 18S nuclear locus. Accession codes and coordinates for the sequences are provided in Supplementary Table S7.

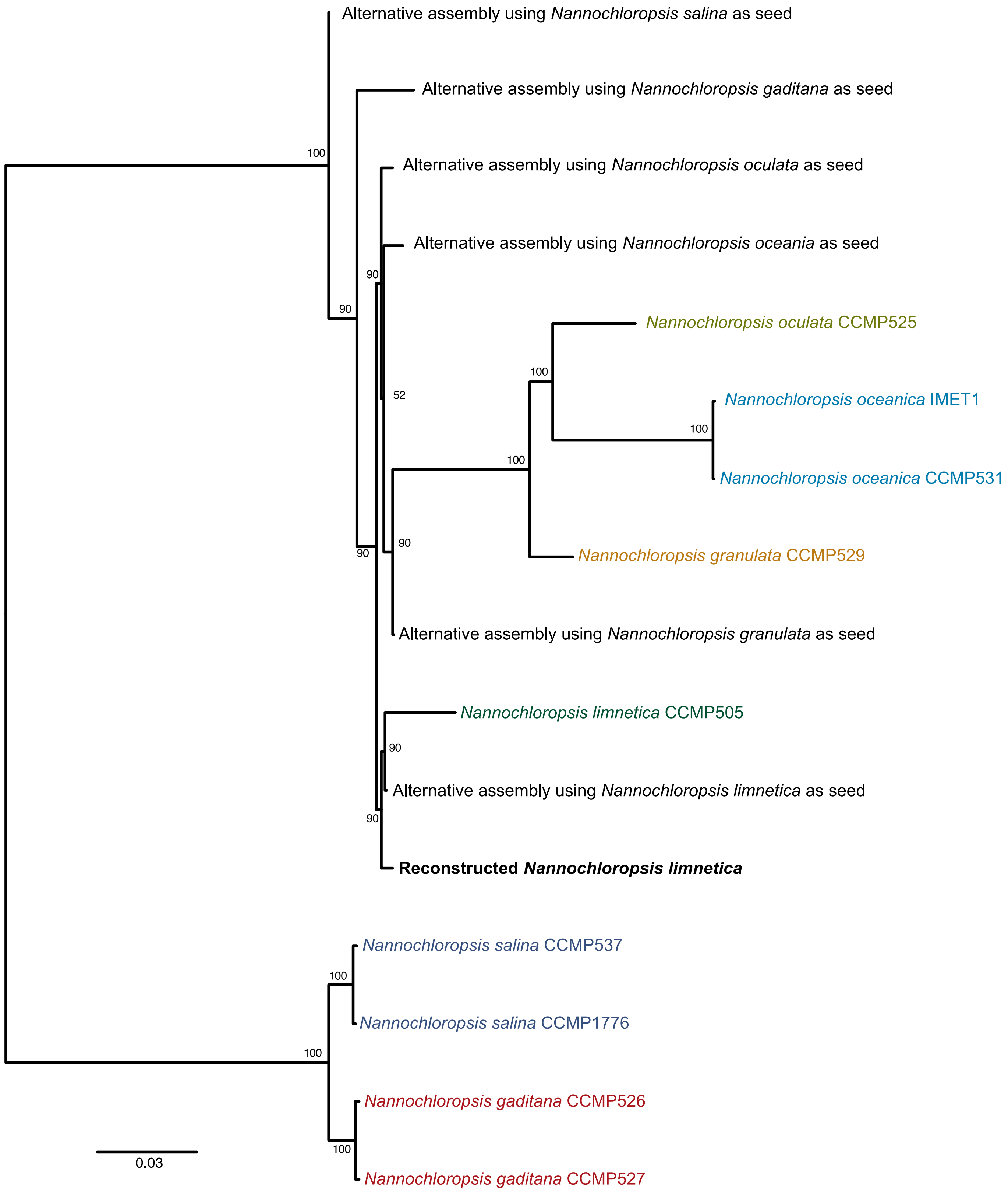


**Supplementary Figure S10**: Maximum likelihood phylogeny of *Nannochloropsis* chloroplast genome sequences, including the reconstructed *N. limnetica* palaeogenomes and the organellar genomes that used alternative *Nannochloropsis* taxa as seed sequences.

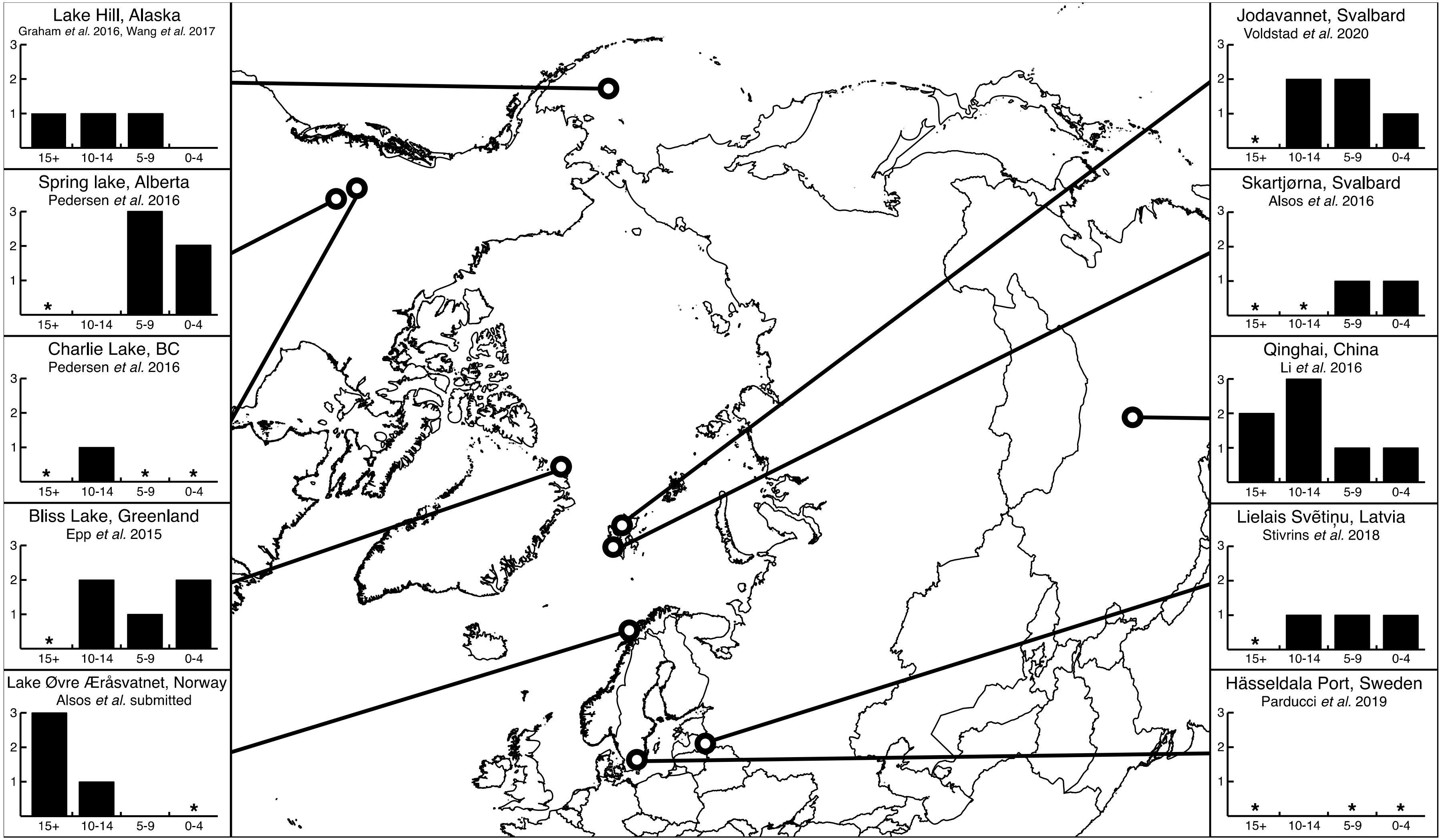


**Supplementary Figure S9**: Overview of *Nannochloropsis* detections, using 10 *seda*DNA data sets either as previously published or based on reanalysis of available data. Plotted on the Y-axis is the inferred abundance of *Nannochloropsis*; 3: dominant in the period, 2: common, 1: rare and 0 is absent. Data was binned into 5000 year time periods, which might obscure finer patterns. No *seda*DNA data was available for time periods marked with an asterisk. References in the main text for: Lake Hill, St. Paul Island, Alaska [1,2], Spring Lake and Charlie Lake, Alberta, Canada [3], Bliss Lake, Greenland [4], Lake Øvre Æråsvatnet, Norway [5], Lakes Jodavannet and Skartjørna, Svalbard [6,7], Lake Qinghai, China [8], Lake Lielais Svētiņu, Latvia [9], and Hässeldala Port, Sweden [10].

**Supplementary Table S1**: The *Nannochloropsis* taxa and the available reference sequences, indicated by an asterisk.

| Species | *rbcL* | 18S | ITS | Chloroplast | Mitochondria | Nuclear genome |
|---|---|---|---|---|---|---|
| *Nannochloropsis australis* | * | * | | | | |
| *Nannochloropsis gaditana* | * | * | * | * | * | * |
| *Nannochloropsis granulata* | * | * | * | * | * | * |
| *Nannochloropsis limnetica* | * | * | * | * | * | * |
| *Nannochloropsis limnetica var. distrophica* | * | * | | | | |
| *Nannochloropsis limnetica var. globosa* | * | * | | | | |
| *Nannochloropsis limnetica var. gutta* | * | | | | | |
| *Nannochloropsis limnetica var. irregularis* | * | | | | | |
| *Nannochloropsis limnetica var. limnetica* | * | * | * | | | |
| *Nannochloropsis maritima* | * | * | * | | | |
| *Nannochloropsis oceanica* | * | * | * | * | * | * |
| *Nannochloropsis oculata* | * | * | * | * | * | * |
| *Nannochloropsis salina* | * | * | * | * | * | * |

**Supplementary Table S2**: Summary statistics for the raw and filtered sequence read counts.

| | **Andøya_LGM_A** | **Andøya_LGM_B** |
|---|---|---|
| Raw paired-end reads | 223,999,748 | 133,265,048 |
| Merged reads | 171,887,513 | 75,784,587 |
| Average read length | 51.44 | 47.9 |
| Percent merged | 76.74 | 56.87 |
| Filtered reads (length >= 35bp and max dust score of 1) | 127,429,489 | 53,000,389 |
| Percent post filtering | 74.14 | 69.94 |
| Average filtered read length | 52.6 | 49.74 |

**Supplementary Table S6**: Summary statistics of the reconstructed *N. limnetica* organellar genomes, as well as the features on the *N. limnetica* reference.

| | **Reconstructed chloroplast** | **Reference chloroplast (NC_022262.1)** | **Reconstructed mitochondria** | **Reference mitochondria (NC_022256.1)** |
|---|---|---|---|---|
| Length | 117,734 | 117,806 | 38,534 | 38,543 |
| Coverage based on the merged Andøya data | 64.3x | 60.8x | 64.9x | 62.4x |
| GC content | 33.34 | 33.54 | 31.68 | 31.69 |
| Coding DNA sequences | 125 | 126 | 35 | 35 |
| Transfer and ribosomal RNAs | 34 | 34 | 28 | 28 |

**Supplementary Table S8**: Reference sequences used as seed sequences for the alternative chloroplast assembly. In addition, the reconstructed length, number of gaps, and the gap length is given for the alternative reconstructed chloroplasts.

| Accession | Reference | Sequence length | Total gap length | Number of gaps | Ungapped length | % Ungapped |
|---|---|---|---|---|---|---|
| KJ410682.1 | *Nannochloropsis gaditana* strain CCMP526 chloroplast, complete genome | 114,710 | 29499 | 1005 | 85,211 | 74.28 |
| KC598085.1 | *Nannochloropsis granulata* chloroplast, complete genome | 117,672 | 3875 | 1615 | 113,797 | 96.7 |
| NC_022262.1 | *Nannochloropsis limnetica* strain CCMP505 chloroplast, complete genome | 117,806 | 2159 | 1695 | 115,647 | 98.17 |
| NC_022263.1 | *Nannochloropsis oceanica* strain CCMP531 chloroplast, complete genome | 117,556 | 5220 | 1548 | 112,336 | 95.56 |
| KC598087.1 | *Nannochloropsis oculata* strain CCMP525 chloroplast, complete genome | 117,462 | 2654 | 1575 | 114,808 | 97.74 |
| NC_022261.1 | *Nannochloropsis salina* strain CCMP537 chloroplast, complete genome | 114,735 | 17,395 | 1266 | 97,340 | 84.84 |

**Supplementary Table S9**: The average number of transversion-only variants for each sample and organellar genome, as well as the average proportion of alternative alleles.

| Sample | Reference | Average number of variants | SD number of variants | Average Variant Ratio | SD Variant Ratio |
|---|---|---|---|---|---|
| Andøya_LGM_A | chloroplast | 376.4 | 4.1593269 | 0.393 | 0.116 |
| Andøya_LGM_B | chloroplast | 299.2 | 2.2803509 | 0.419 | 0.122 |
| Andøya_LGM_A | mitochondria | 111.8 | 0.4472136 | 0.385 | 0.108 |
| Andøya_LGM_B | mitochondria | 80.6 | 0.8944272 | 0.427 | 0.108 |

**Supplementary Table S11**: The *Nannochloropsis* occurrences detected in a contemporary northern Norway environmental DNA metabarcoding data set [11].

| Sample | Lake | District | Habitat type | Core | Depth | *Nannochloropsis* reads | *Nannochloropsis* PCR replicates (out of 6) |
|---|---|---|---|---|---|---|---|
| TTE01 | Finnvatnet | Kvaløya | Birch forest/mire | Core1 | 0-2cm | 0 | 0 |
| TTE03 | Lakselvhøgda | Ringvassøya | Alpine heath and mire | Core1 | 0-2cm | 0 | 0 |
| TTE04 | Lakselvhøgda | Ringvassøya | Alpine heath and mire | Core1 | 2-4cm | 0 | 0 |
| TTE05 | Lakselvhøgda | Ringvassøya | Alpine heath and mire | Core2 | 0-2cm | 0 | 0 |
| TTE06 | Jula Jávri | Kåfjorddalen | Alpine heath and mire | Core1 | 0-2cm | 736 | 5 |
| TTE07 | Jula Jávri | Kåfjorddalen | Alpine heath and mire | Core2 | 0-2cm | 107 | 4 |
| TTE09 | Lauvås | Ringvassøya | Heath, mire and birch forest | Core1 | 0-2cm | 0 | 0 |
| TTE10 | Lauvås | Ringvassøya | Heath, mire and birch forest | Core1 | 2-4cm | 0 | 0 |
| TTE11 | Lauvås | Ringvassøya | Heath, mire and birch forest | Core2 | 0-2cm | 0 | 0 |
| TTE12 | Øvre Æråsvatnet | Andøya | Mires and birch forest, conifers planted | Core1 | 0-2cm | 0 | 0 |
| TTE13 | Øvre Æråsvatnet | Andøya | Mires and birch forest, conifers planted | Core2 | 0-2cm | 0 | 0 |
| TTE14 | Paulan Jávri | Kåfjorddalen | Alpine heath | Core1 | 0-2cm | 0 | 0 |
| TTE15 | Paulan Jávri | Kåfjorddalen | Alpine heath | Core2 | 0-2cm | 0 | 0 |
| TTE17 | Gauptjern | Dividalen | Sub-alpine birch forest | Core1 | 0-2cm | 0 | 0 |
| TTE18 | Gauptjern | Dividalen | Sub-alpine birch forest | Core1 | 2-4cm | 26 | 1 |
| TTE19 | Gauptjern | Dividalen | Sub-alpine birch forest | Core1 | 4-6cm | 0 | 0 |
| TTE20 | Gauptjern | Dividalen | Sub-alpine birch forest | Core2 | 0-2cm | 0 | 0 |
| TTE21 | Gauptjern | Dividalen | Sub-alpine birch forest | Core2 | 2-4cm | 0 | 0 |
| TTE22 | Gauptjern | Dividalen | Sub-alpine birch forest | Core2 | 4-6cm | 0 | 0 |
| TTE23 | Brennskogtjønna | Dividalen | Pine forest | Core1 | 0-2cm | 86 | 2 |
| TTE25 | Brennskogtjønna | Dividalen | Pine forest | Core1 | 2-4cm | 0 | 0 |
| TTE26 | Brennskogtjønna | Dividalen | Pine forest | Core1 | 4-6cm | 217 | 4 |
| TTE27 | Brennskogtjønna | Dividalen | Pine forest | Core2 | 0-2cm | 358 | 3 |
| TTE28 | Brennskogtjønna | Dividalen | Pine forest | Core2 | 2-4cm | 127 | 1 |
| TTE29 | Brennskogtjønna | Dividalen | Pine forest | Core2 | 4-6cm | 179 | 1 |
| TTE30 | A-tjern | Dividalen | Birch forest/mire | Core1 | 0-2cm | 3732 | 6 |
| TTE31 | A-tjern | Dividalen | Birch forest/mire | Core1 | 2-4cm | 4317 | 6 |
| TTE33 | A-tjern | Dividalen | Birch forest/mire | Core1 | 4-6cm | 4073 | 6 |
| TTE34 | A-tjern | Dividalen | Birch forest/mire | Core2 | 0-2cm | 4528 | 6 |
| TTE35 | A-tjern | Dividalen | Birch forest/mire | Core2 | 2-4cm | 7314 | 6 |
| TTE36 | A-tjern | Dividalen | Birch forest/mire | Core2 | 4-6cm | 12899 | 6 |
| TTE37 | Rottjern | Dividalen | Pine and birch forest | Core1 | 0-2cm | 5259 | 6 |
| TTE38 | Rottjern | Dividalen | Pine and birch forest | Core1 | 2-4cm | 6092 | 6 |
| TTE39 | Rottjern | Dividalen | Pine and birch forest | Core1 | 4-6cm | 4435 | 6 |
| TTE41 | Rottjern | Dividalen | Pine and birch forest | Core1 | 6-8cm | 2273 | 6 |
| TTE42 | Rottjern | Dividalen | Pine and birch forest | Core2 | 0-2cm | 2973 | 6 |
| TTE43 | Rottjern | Dividalen | Pine and birch forest | Core2 | 2-4cm | 4170 | 6 |
| TTE44 | Rottjern | Dividalen | Pine and birch forest | Core2 | 4-6cm | 2130 | 6 |
| TTE45 | Rottjern | Dividalen | Pine and birch forest | Core2 | 6-8cm | 3565 | 6 |
| TTE46 | Einletvatnet | Andøya | Mires, patches of birch forest | Core1 | 0-2cm | 0 | 0 |
| TTE47 | Einletvatnet | Andøya | Mires, patches of birch forest | Core2 | 0-2cm | 0 | 0 |

**Supplementary Table S12**: Accession codes used for the annotation of the reconstructed organellar genomes.

| Species | Chloroplast accession | Mitochondrial accession |
|---|---|---|
| *Nannochloropsis gaditana* | NC_020014.1 | NC_020015.1 |
| *Nannochloropsis granulata* | NC_022259.1 | NC_022254.1 |
| *Nannochloropsis limnetica* | NC_022262.1 | NC_022256.1 |
| *Nannochloropsis oceanica* | NC_022263.1 | NC_022258.1 |
| *Nannochloropsis oculata* | NC_022260.1 | NC_022257.1 |

### External Supplementary Tables

**Supplementary Table S3:** Tabulated MEGAN output for both samples and the non-overlapping one million read subsets.

**Supplementary Table S4:** The reference genomes used for the mapping analysis along with the raw and filtered sequence counts, corrected sequence counts and coverage for each sample. In addition, the shared and unique sequence counts are provided between *Nannochloropsis* nuclear and chloroplast genomes, as well as the *Mycobacterium* genomes.

**Supplementary Table S5:** Counts of sequences across the different LCA levels for the organellar genome mapping analysis. In addition, lists of accession codes are provided for the organellar genomes used as reference.

**Supplementary Table S7:** Accession codes for the *Nannochloropsis* sequences used in the organellar and *rbcL*, 18S and ITS phylogenies. For the smaller markers the start and end coordinates used for phylogenetic analysis are given for each accession.

**Supplementary Table S10:** The raw linked allele analysis results for both samples and organellar genomes as well as the summarized results.

**Supplementary Table S13:** Detections of N. limnetica detections in the re-analysed Lake Hill, St. Paul Island, Alaska [1,2], Spring Lake and Charlie Lake, Alberta, Canada [3] and Hässeldala Port, Sweden [10] shotgun sequencing datasets.

**Supplementary Table S14:** The sample names and PCR tags used for the reanalysed Skartjørna, Svalbard, sedaDNA metabarcode dataset [6].
